# *C. elegans* XMAP215/ZYG-9 and TACC/TAC-1 act at multiple times throughout oocyte meiotic spindle assembly to promote both the coalescence of pole foci into a bipolar structure, and the stability of coalesced poles, during oocyte meiotic cell division

**DOI:** 10.1101/2022.08.02.502432

**Authors:** Austin M. Harvey, Bruce Bowerman

## Abstract

The conserved two-component XMAP215/TACC modulator of microtubule stability is required in multiple animal phyla for acentrosomal spindle assembly during oocyte meiotic cell division, with *C. elegans XMAP215/zyg-9* and *TACC/tac-1* mutant oocytes exhibiting multiple and indistinguishable defects beginning early in meiosis I. To determine if these defects represent one or more early requirements with additional later and indirect consequences, or multiple temporally distinct and more direct requirements, we have used live cell imaging and fast-acting temperature-sensitive *zyg-9* and *tac-1* alleles to dissect at high temporal resolution their oocyte meiotic spindle assembly requirements. Our results from temperature upshift and downshift experiments indicate that the ZYG-9/TAC-1 complex has multiple temporally distinct and separable requirements throughout oocyte meiotic cell division. First, we show that during prometaphase ZYG-9 and TAC-1 promote the coalescence of early pole foci into a bipolar structure both by stabilizing pole foci as they grow and by limiting their growth rate, with these requirements being independent of an earlier defect in microtubule organization. Second, during metaphase, ZYG-9 and TAC-1 maintain spindle bipolarity by suppressing ectopic pole formation, and this pole stability is important for maintaining chromosome congression. Finally, we show that ZYG-9 and TAC-1 also are required for the proper coalescence of pole foci during meiosis II, independently of their requirements during meiosis I. Our findings highlight the value of fast-acting temperature-sensitive alleles for high resolution temporal dissection of gene requirements, and we discuss how negative regulation of microtubule stability by ZYG-9/TAC-1 during oocyte meiotic cell division might account for the observed defects in spindle pole coalescence and stability.

**Author Summary:** When most animal cells divide, organizing centers called centrosomes nucleate and organize protein filaments called microtubules into a dynamic bipolar structure called the spindle that equally partitions the duplicated genome between two daughter cells. However, female egg cells, called oocytes, lack centrosomes but still assemble bipolar spindles that separate chromosomes. Using the nematode *C. elegans* as a model system, and taking advantage of fast-acting temperature-sensitive alleles of both ZYG-9 and TAC-1 that rapidly inactivate or reactivate upon temperature upshifts or downshifts, respectively, we show that a complex of two regulators of microtubule stability, called ZYG-9 and TAC-1, has multiple and separable requirements during the process of acentrosomal oocyte spindle assembly. These requirements include promoting the coalescence of early pole foci into a bipolar structure, and the maintenance of pole stability after the assembly of a bipolar structure, both of which are essential for proper chromosome separation. Furthermore, oocyte meiosis involves two consecutive cell divisions, and we show that ZYG-9/TAC-1 are required for pole coalescence during both the first and second meiotic cell divisions. Our findings provide a high resolution view of the distinct and separable temporal requirements for these widely conserved regulators of microtubule stability during acentrosomal oocyte spindle assembly.

## Introduction

Oocyte meiosis I and II are sequential, highly asymmetric cell divisions that reduce a duplicated genome to single copy, producing a haploid egg. Meiotic cell division requires a microtubule-based spindle apparatus to partition the genome and, in contrast to mitotic cells, the oocytes of most animal models lack centrosomes but nevertheless nucleate and organize microtubules into functional bipolar spindles (Dumont and Desai, 2012; Severson et al., 2016; Mullen et al., 2019; Ohkura, 2015). Faithful transmission of the genome is essential, but the molecular mechanisms underlying acentrosomal oocyte meiotic spindle assembly and function *in vivo* remain poorly understood.

In *Caenorhabditis elegans*, oocyte spindle assembly occurs through a sequence of stages defined by visible changes in microtubule dynamics (Gigant et al., 2017; Wolff et al., 2016). First is nuclear envelope breakdown (NEBD) and the entry of α- and β-tubulin heterodimers into the nucleus. Microtubule bundles then appear beneath the disassembling nuclear lamina, forming a peripheral and roughly spherical microtubule network at the cage stage. Next, during the multipolar stage, microtubules increase in amount and assemble into a network with numerous nascent pole foci marked with the pole protein ASPM-1 (Connolly et al., 2015). Over time these pole foci coalesce, forming a bipolar spindle by metaphase, with homologous chromosome pairs aligned between the poles.

Following these stages in bipolar spindle assembly, the oocyte transitions into anaphase, when the spindle undergoes extensive morphological changes during chromosome separation (Yang et al., 2003, 2005). First, the spindle shortens and the poles broaden, as the spindle rotates to orient perpendicularly to the cortex (McNally et al., 2016), with the two separating chromosome sets each moving slightly towards the nearest pole during anaphase A. Subsequently, the spindle poles largely disassemble while parallel arrays of microtubules appear and elongate between the separating chromosome sets during anaphase B (Laband et al., 2017; Davis-Roca et al., 2018; Danlasky et al., 2020), with half of the genome extruded into the first polar body upon the completion of meiosis I. While this description provides a platform for further investigation, when and how the proteins required for oocyte spindle assembly function during these different stages and transitions remains poorly understood.

XMAP215 and the Transforming and Acidic Coiled-Coil (TACC) are widely conserved proteins, with family members that can bind each other and often function as a complex to regulate microtubule stability in a variety of cellular contexts, including *C. elegans* oocyte meiotic cell division (Le Bot et al., 2003; Srayko et al., 2005; Bellanger and Gönczy, 2003; Peset and Vernos, 2008; Al-Bassam and Chang, 2011; Chen et al., 2015; Chuang et al., 2020; So et al., 2019). XMAP215 orthologs have a C-terminal acidic domain that binds microtubules and multiple TOG (Tumor Over-expressed Gene) domains that bind to and increase the local concentration of tubulin heterodimers, modulating their incorporation or removal at microtubule plus ends (Al-Bassam et al., 2007; Akhmanova and Steinmetz, 2015). Studies in *Xenopus* and in *Drosophila* have identified a C-terminal TACC domain that mediates binding to XMAP215 proteins and promotes the localization of both XMAP215 and TACC proteins to centrosomes (Gergely et al., 2000; Lee et al., 2001; Peset et al., 2005). During the first mitotic division of the early *C. elegans* embryo, XMAP215/ZYG-9 and TACC/TAC-1 co-localize to centrosomes and spindle microtubules, mutually depend on each other for protein stability, and are both required for spindle microtubule stability and proper mitotic spindle positioning (Bellanger and Gönczy, 2003; Srayko et al., 2003; Bellanger et al., 2007).

While the mitotic functions of ZYG-9 and TAC-1 have been investigated more extensively, their roles during *C. elegans* oocyte meiotic cell division are not as well understood. Early studies showed that ZYG-9 is concentrated at oocyte spindle poles and diffusely associated with spindle microtubules (Matthews et al., 1998), and RNA interference (RNAi) knockdown of ZYG-9 resulted in disorganized spindles and frequent chromosome separation errors (Matthews et al., 1998; Yang et al., 2003). More recently, live imaging studies have shown that ZYG-9 and TAC-1 exhibit indistinguishable spindle localization patterns during oocyte meiotic cell division, and RNAi depletion of either protein resulted in identical phenotypes (Chuang et al., 2020). The earliest detected defects were observed at the microtubule cage stage, when microtubule bundles were not restricted to the periphery but sometimes passed through the internal chromosome-occupied space. Subsequently the coalescence of pole foci was defective, with foci sometimes splitting, and mutant oocytes sometimes forming tripolar spindles that separated chromosomes into three sets.

Notably, in contrast to their well-documented role in promoting microtubule stability during early embryonic mitosis, ZYG-9 and TAC-1 negatively regulate microtubule stability during oocyte meiotic cell division. RNAi knockdown of either results in significantly elevated microtubule levels in oocytes, both in association with the egg chromosomes and throughout the cortex (Chuang et al., 2020). While genetic studies in budding yeast (van Breugel et al., 2003; Kosco et al., 2001) and in *Drosophila* S2 cells (Brittle and Ohkura, 2005), and biochemical studies in *Xenopous* egg extracts (Shirasu-Hiza et al., 2003), have also suggested that XMAP215 orthologs can destabilize microtubules in some contexts, most studies have focused on their roles in promoting microtubule stability and growth (Akhmanova and Steinmetz, 2015; Cook et al., 2019; So et al., 2019). *C. elegans* oocyte meiotic cell division thus provides an appealing model for investigating the negative regulation of microtubule stability by XMAP215 and TACC family members.

The numerous defects observed upon loss of ZYG-9 or TAC-1 suggest that this complex may have multiple separate requirements during oocyte meiotic cell division. However, the use of non-conditional mutations or knockdown methods to assess gene requirements precludes determining whether each defect represents a separate requirement, or if instead the later defects are indirect consequences of earlier ones. The use of CRISPR/Cas9 genome editing to degron-tag genes for auxin-inducible degradation now makes it possible to engineer conditional loss of function alleles throughout the genome (Zhang et al., 2015). However, meiosis I in *C. elegans* is rapid, progressing from nuclear envelope breakdown to polar body extrusion over a roughly 20 to 35 minute interval (Chuang et al., 2020), limiting the utility of degron-tagging for dissecting gene requirements with high temporal resolution.

Temperature-sensitive (TS) alleles, typically missense mutations that produce functional proteins at lower permissive temperatures but are inactive at higher restrictive temperatures, provide an alternative and powerful approach for the temporal dissection of gene requirements. Many *C. elegans* TS alleles can be classified as either slow- or fast-acting (O’Rourke et al., 2011). With slow-acting alleles, fertile worms must be cultured for hours at the restrictive temperature to observe mutant phenotypes. Such slow-acting alleles are likely due to mutations that cause irreversible protein folding outcomes and require the production of newly synthesized proteins and oocytes for the consequences of a temperature upshift or downshift to be observed. By contrast, fast-acting, heat-sensitive TS alleles rapidly inactivate within a few minutes of a temperature upshift, even if the proteins were translated at a lower permissive temperature, likely because the missense mutations make the proteins less thermally stable and more prone to unfolding or inactivation at the higher restrictive temperatures. Some fast-acting TS alleles also are reversible, with protein function rapidly restored upon shifting from restrictive to permissive temperatures, adding to their usefulness (Severson et al., 2000). To better define the requirements for ZYG-9 and TAC-1 during oocyte meiotic cell division, we have taken advantage of fast-acting TS alleles, with live imaging and fluorescent protein fusions, to identify multiple and separate stage-specific meiotic requirements for these two widely conserved regulators of microtubule dynamics.

## Results

### Temperature-sensitive *XMAP215/zyg-9* and *TACC/tac-1* alleles are fast-acting during *C. elegans* oocyte meiotic cell division

We have employed three previously isolated recessive and temperature-sensitive alleles—*zyg-9(or623ts), zyg-9(or634ts)*, and *tac-1(or455ts)*—to temporally dissect these gene requirements during *C. elegans* oocyte meiotic spindle assembly (Bellanger et al., 2007).

Hereafter, we collectively refer to the mutant oocytes as *zyg-9* and *tac-1* mutants, or as TS mutants. These alleles all showed less than 1% embryonic-lethality when homozygous mutant adults were cultured at the permissive temperature of 15°C and greater than 99% embryonic-lethality when cultured at the restrictive temperature of 26°C (Bellanger et al., 2007), and therefore were likely to be fast-acting (O’Rourke et al., 2011). To observe spindle assembly dynamics, we used spinning disk confocal microscopy, coupled with a microfluidics temperature-control system, to image live oocytes *in utero* within immobilized whole-mount worms from transgenic strains that express both a GFP-tagged ß-tubulin (GFP::TBB-2) to mark microtubules, and an mCherry-tagged histone (mCherry::H2B) to mark chromosomes (see Materials and Methods).

First, we compared oocyte meiotic spindle assembly in control and mutant oocytes maintained at 15°C throughout meiosis I, to determine if any non-essential defects might occur even when mutant oocytes were kept at the permissive temperature. When maintained at 15°C, *zyg-9* and *tac-1* mutant oocytes routinely formed barrel-shaped bipolar spindles that aligned chromosomes at the spindle midpoint, with spindle assembly and chromosome separation dynamics indistinguishable from those observed in control oocytes, except for lagging chromosomes during anaphase that we saw in 2 of 20 *zyg-9(or623ts)* and 1 of 13 *zyg-9(or634ts)* oocytes maintained at 15°C (Fig. 1A-D, Fig. S1). These results indicate that any further defects observed after temperature upshifts are due to inactivation of the mutant protein.

**FIGURE 1.**
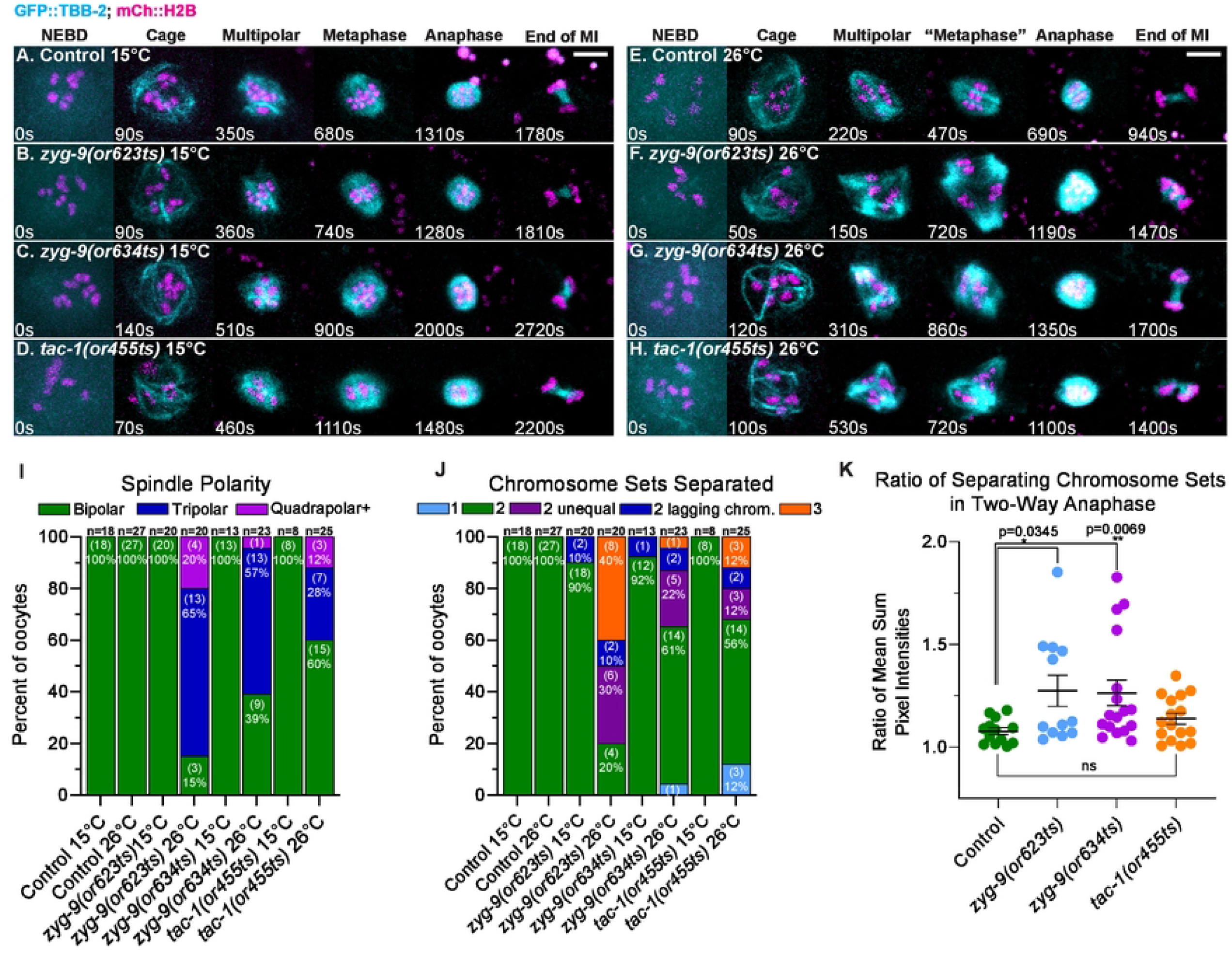
Oocyte meiosis I spindle assembly dynamics in *zyg-9* and *tac-1* TS mutants at the permissive and restrictive temperatures. (A-H) Time-lapse maximum intensity projection images during meiosis I in live control and TS mutant oocytes expressing GFP::TBB-2 and mCherry::H2B to mark microtubules and chromosomes, at 15°C (A-D) and at 26°C (E-H). In this and in all subsequent meiosis I time-lapse image series, t=0 is labeled NEBD and is the timepoint immediately preceding the appearance of microtubule bundles forming the cage structure. To account for differences in movie signal quality inherent to *in utero* live cell imaging, the intensity scales for montages in all figures were set individually to give the clearest depiction of the spindle and chromosomes. (see Materials and Methods). (I-J) Number of spindle poles present at the onset of spindle shortening per oocyte (I), and number of chromosome sets separated after anaphase per oocyte (J); number of oocytes examined indicated above each bar, number scored with each phenotype shown in parentheses inside bars with corresponding percent below. (K) Ratio of fluorescence intensity of two separated chromosome sets at 26°C (see 1J) at the end of anaphase B. For all figures, distributions of scatter plot values were compared using the Mann–Whitney U-test to calculate P-values. Error bars and values are mean ± SEM. *, P <0.05; **, P < 0.01. Scale bars = 5 μm. See the Materials and Methods for a description of the spindle assembly stages and frame selection in this and all other figures.

We next assessed how effectively these TS alleles reduce gene function. We compared the defects observed during spindle assembly when TS mutant oocytes were maintained at the restrictive temperature throughout meiosis I, to the defects previously reported following strong RNAi knockdowns (Matthews et al., 1998; Yang et al., 2003; Chuang et al., 2020). While control oocytes maintained at 26°C always became bipolar with normal spindle assembly dynamics (Fig. 1E, Movie S1, 18/18 oocytes), spindle microtubules and chromosomes in TS mutant oocytes maintained at 26°C were highly disorganized, with spindles that frequently failed to become bipolar (Fig. 1F-H, Fig. S1, Movie S2), consistent with previous studies using RNAi to knockdown ZYG-9 and TAC-1 (Matthews et al., 1998; Chuang et al., 2020).

To more quantitatively compare assembly dynamics and spindle polarity in these different genetic backgrounds, we used the beginning of spindle shortening, which in wild-type oocytes occurs upon the transition from metaphase to anaphase (Yang et al., 2003, 2005). Even in highly abnormal *zyg-9* or *tac-1* mutant spindles that formed at 26°C, spindle shortening was easily identifiable (Fig. 1F-H, Fig. S1, Movie S2), as reported previously after RNAi knockdown (Yang et al., 2003). When maintained at 26°C, spindle bipolarity was established by the beginning of spindle shortening in only 3 of 20 *zyg-9(or623ts)*, 9 of 23 *zyg-9(or634ts)*, and 15 of 25 *tac-1(or455ts)* oocytes, whereas in mutant oocytes kept at 15°C, all spindles were bipolar by the beginning of spindle shortening (Fig. 1I).

We also observed severe chromosome separation errors in *zyg-9* and *tac-1* mutant oocytes maintained at the restrictive temperature throughout meiosis I. By late in meiosis I, we often observed separation outcomes that resulted in one or three, instead of two, chromosome sets, as well as lagging chromosomes bridging the sets (Fig. 1J). While separation into two sets occurred in roughly half of the TS mutant oocytes, the chromosomal distribution between the two sets was often unequal (Fig. 1K), whereas we consistently observed two equally sized chromosome sets in TS mutant oocytes maintained at 15°C, as in control oocytes (Fig. 1J, 1K).

The penetrance of the severe separation defects, into one or three sets, roughly matched the penetrance of the failures to establish bipolar spindles, suggesting that the defects in spindle assembly may account for these subsequent errors in chromosome separation. However, while roughly half of mutant oocytes separated chromosomes into three sets after ZYG-9 RNAi knockdown (Chuang et al., 2020), we observed fewer examples (∼1/3 overall) of three-way separation in TS mutant oocytes at the restrictive temperature (Fig. 1J). We conclude that *zyg-9* and *tac-1* mutant oocytes (i) assemble spindles with roughly normal dynamics at the permissive temperature, and (ii) exhibit spindle assembly and polarity defects at the restrictive temperature very similar to those previously observed after RNAi knockdown, albeit with somewhat lower penetrance.

Finally, we used temperature upshifts to determine if these TS *zyg-9* and *tac-1* alleles are fast-acting. To do so, we upshifted mutant oocytes from the permissive temperature of 15°C to the restrictive temperature of 26°C at the multipolar stage, between roughly 2.7 and 6.8 minutes after cage onset, preceding the establishment of spindle bipolarity (Materials and Methods, Fig. S2). We then observed the subsequent spindle assembly and chromosome separation dynamics. Prior to the upshift, microtubule networks in *zyg-9* and *tac-1* oocytes were coalescing into a bipolar structure as in control oocytes (Fig. 2B-D, Fig. S3). Immediately after the upshifts, coalescence was disrupted, and the penetrance of the subsequent spindle assembly and chromosome separation defects were nearly identical to those observed in TS mutant oocytes maintained at 26°C throughout meiosis I (Figs. 1I, 1J, 2I, 2J, Movies S3-S6). In contrast, temperature upshifts at the multipolar stage in control oocytes had no effect on pole coalescence, spindle bipolarity and chromosome separation (Fig. 2A; 7/7 oocytes, Fig. S3). We conclude that *zyg-9(or623ts), zyg-9(or634ts)*, and *tac-1(or455ts)* are fast-acting during oocyte meiotic cell division.

**FIGURE 2.**
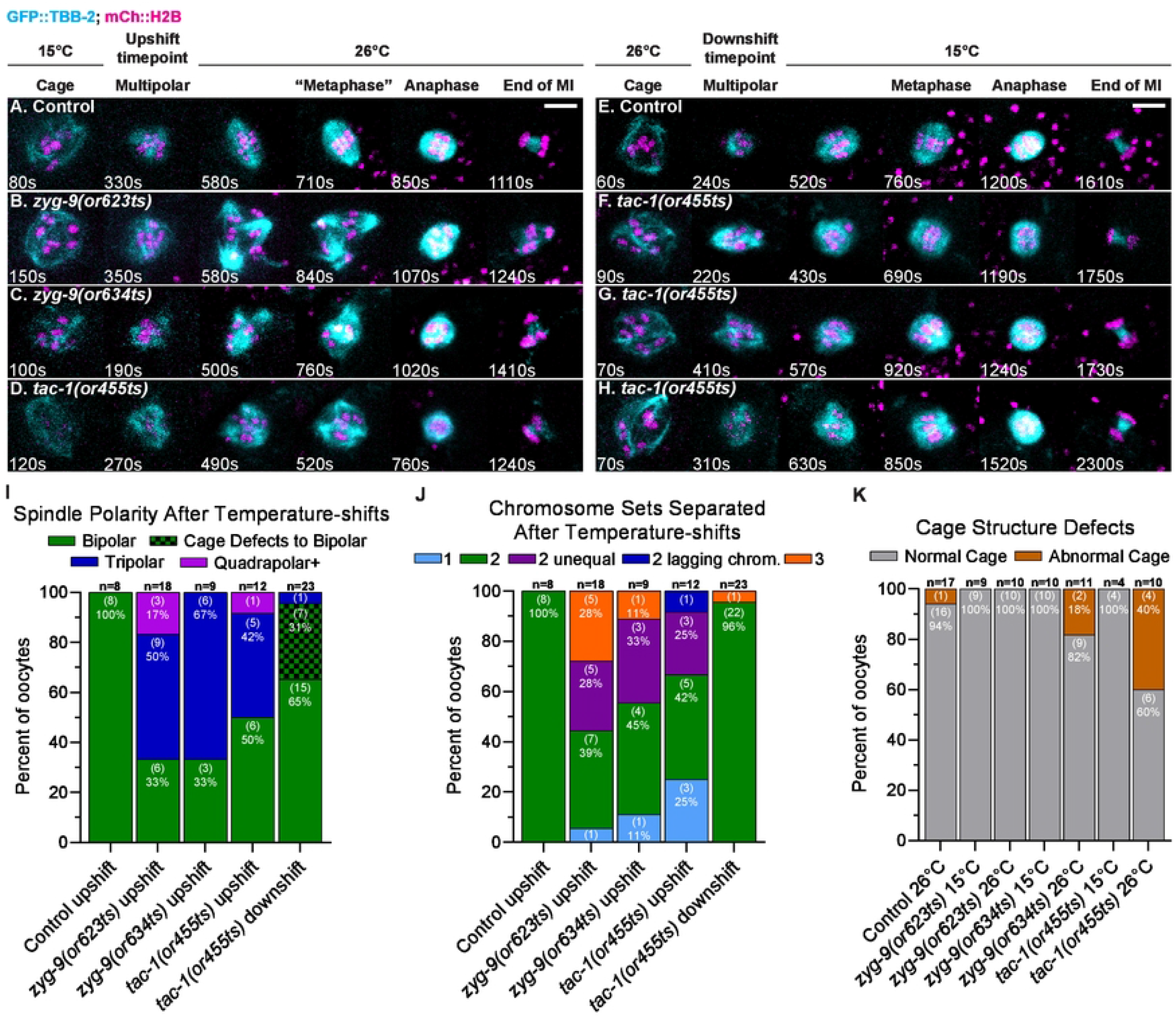
Pole coalescence and polarity defects in *zyg-9* and *tac-1* mutant oocytes do not depend on prior cage structure defects. (A-H) Time-lapse maximum intensity projection images of live control and TS mutant oocytes upshifted from 15°C to 26°C (A-D) or downshifted from 26°C to 15°C (E-H) during the multipolar stage, in oocytes expressing GFP::TBB-2 and mCherry::H2B. Oocytes depicted in F-H have cage structure defects (for an example in 3-D, see Movie S9). (I-K) Number of spindle poles present at the onset of spindle shortening per oocyte (I); number of chromosome sets separated during anaphase per oocyte (J); number of oocytes with abnormal cage structures (K); number of oocytes examined indicated above each bar, number scored with each phenotype shown in parentheses inside bars with corresponding percent below. Bar graphs in (I) and (J) include oocytes expressing GFP::TBB-2 (this figure) and oocytes expressing GFP::ASPM-1 (see Figure 4).

### Spindle polarity and chromosome separation defects in *zyg-9* and *tac-1* mutants do not depend on prior cage structure defects

Having established that these TS alleles are fast-acting, we then asked whether the later spindle bipolarity and chromosome separation defects in *zyg-9* and *tac-1* mutant oocytes depend on the abnormal microtubule cage structure that forms shortly after NEBD, the earliest defect described thus far in these mutants. In some *zyg-9* and *tac-1* mutant oocytes, the microtubule bundles that form the cage are not restricted to the periphery as in control oocytes (Movie S7), but also pass through the chromosome occupied space inside the cage (Chuang et al., 2020). Because the cage structure might promote the coalescence of early pole foci by making the process two dimensional, rather than three-dimensional throughout the volume occupied by chromosomes, a causal relationship between this cage structure defect and the subsequent pole coalescence defect is appealing (Chuang et al., 2020). However, when we examined *zyg-9* and *tac-1* mutant oocytes that failed to establish a bipolar spindle after multipolar stage temperature upshifts, we found that the cage structures had formed properly prior to the upshifts, with microtubule bundles restricted to the periphery (Fig. 2B-D, Fig. S3, Movie S8). These results suggest that the pole coalescence defects in *zyg-9* and *tac-1* do not depend on an earlier detectably abnormal cage structure.

We next performed complimentary temperature-downshift experiments to ask if bipolar spindle assembly can be rescued in mutant oocytes with earlier cage defects. For these downshift experiments, we used *tac-1(or455ts)*, as it resulted in the most highly penetrant cage structure defect when TS mutants were kept at 26°C throughout meiosis I (Fig. 2K, Fig. S3G). In *tac-1(or455ts)* oocytes kept at 26°C until the multipolar stage, and then downshifted to 15°C, between roughly 2.3 and 6.2 minutes after NEB (Fig. S2), the spindle microtubules coalesced after the downshifts into bipolar spindles that separated chromosomes into two equal sets in 22 of 23 mutant oocytes, with the one exception being a mutant oocyte that separated chromosomes into three sets (Fig. 2F-I). Importantly, bipolar spindles assembled and separated chromosomes into two equal sets in all seven of the mutant oocytes in which we observed early cage structure defects (Fig. 2J, Movie S9), indicating that cage structure defects are not sufficient to cause later coalescence defects. Based on these temperature upshift and downshift experiments, we conclude that the pole coalescence defects observed during prometaphase in *zyg-9* and *tac-1* mutants occur independently of the earlier cage structure defects.

### The spindle polarity and chromosome separation defects in *zyg-9* and *tac-1* mutant oocytes do not depend on elevated microtubule levels

Another mutant phenotype in *zyg-*9 and *tac-1* oocytes that could indirectly cause defects in spindle bipolarity and chromosome separation is the prominent accumulation of abnormally high levels of microtubules, both in association with oocyte chromosomes and also throughout the oocyte cortex during meiosis I spindle assembly, although the levels vary substantially from oocyte to oocyte, and over time in any one oocyte (Chuang et al., 2020). To investigate whether increased spindle microtubule levels might be responsible for the later failures to establish spindle bipolarity and properly separate chromosomes, we quantified spindle-associated microtubule levels in TS mutants oocytes maintained at 26°C throughout meiosis I (Materials and Methods). Surprisingly, spindle microtubules levels were significantly increased only in *tac-1(or455ts)* oocytes but not in *zyg-9(or623ts)* or *zyg-9(or634ts)* oocytes, relative to control oocytes maintained at 26°C (Fig. 3A, 3B). Because spindle assembly and chromosome separation were often defective in all three TS mutants when maintained at 26°C, a simple elevation in overall spindle microtubule levels alone cannot account for the spindle bipolarity and chromosome separation defects observed after reducing ZYG-9 or TAC-1 function (see Discussion).

**Figure 3.**
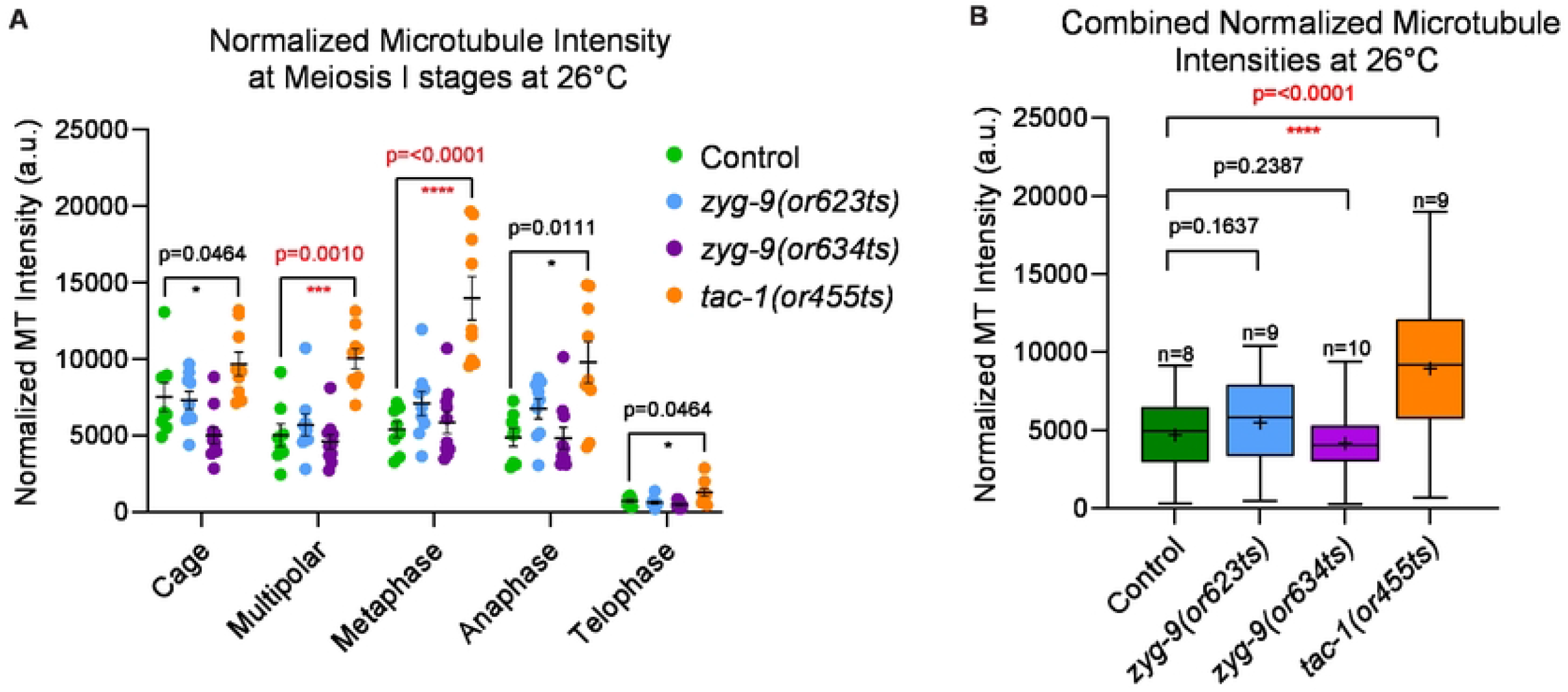
Microtubule levels are elevated throughout meiosis I in *tac-1(or455ts)* oocytes. (A-B) Normalized microtubule pixel intensity at 26°C in arbitrary units (see Materials and Methods) for (A) oocyte spindles at each stage of meiosis I, error bars and values are mean ± SEM; and (B) all meiotic stages combined for control and TS mutant oocytes. The boxplots display the datasets, with the median (line) and mean (+) intensity values for control and mutant oocytes; bars are 75% and whiskers are 95% of the observed normalized microtubule intensity values. *, P <0.05; **, P < 0.01; ***, P < 0.001; ****, P <0.0001. Scale bars = 5 μm.

### ZYG-9 and TAC-1 prevent the splitting of early pole foci and limit their growth during bipolar spindle formation

We next used temperature upshift and downshift experiments to ask how ZYG-9 and TAC-1 influence the prometaphase coalescence of early pole foci. To assess pole coalescence, we used live imaging of control and mutant oocytes from transgenic strains that express an endogenous GFP fusion to the spindle pole marker ASPM-1 (GFP::ASPM-1), and the mCherry::H2B fusion to mark chromosomes (Fig. 4, Figs. S4 & S5). Soon after NEBD in control and mutant oocytes maintained at 15°C throughout meiosis I, diffuse clouds with small GFP::ASPM-1 foci formed a network around the chromosomes, coalescing over time into fewer and larger foci and ultimately forming bipolar spindles with chromosomes aligned midway between the poles (Fig. 4A-C, Fig. S4). In contrast, in mutant but not control oocytes kept at 26°C throughout meiosis I, the early pole foci were more dynamic, often splitting apart instead of coalescing, and in some cases the mutant oocytes ultimately assembled spindles with three poles (Fig. 4D-F, Fig. S4), consistent with the defects previously reported after RNAi knockdowns (Chuang et al., 2020).

**FIGURE 4.**
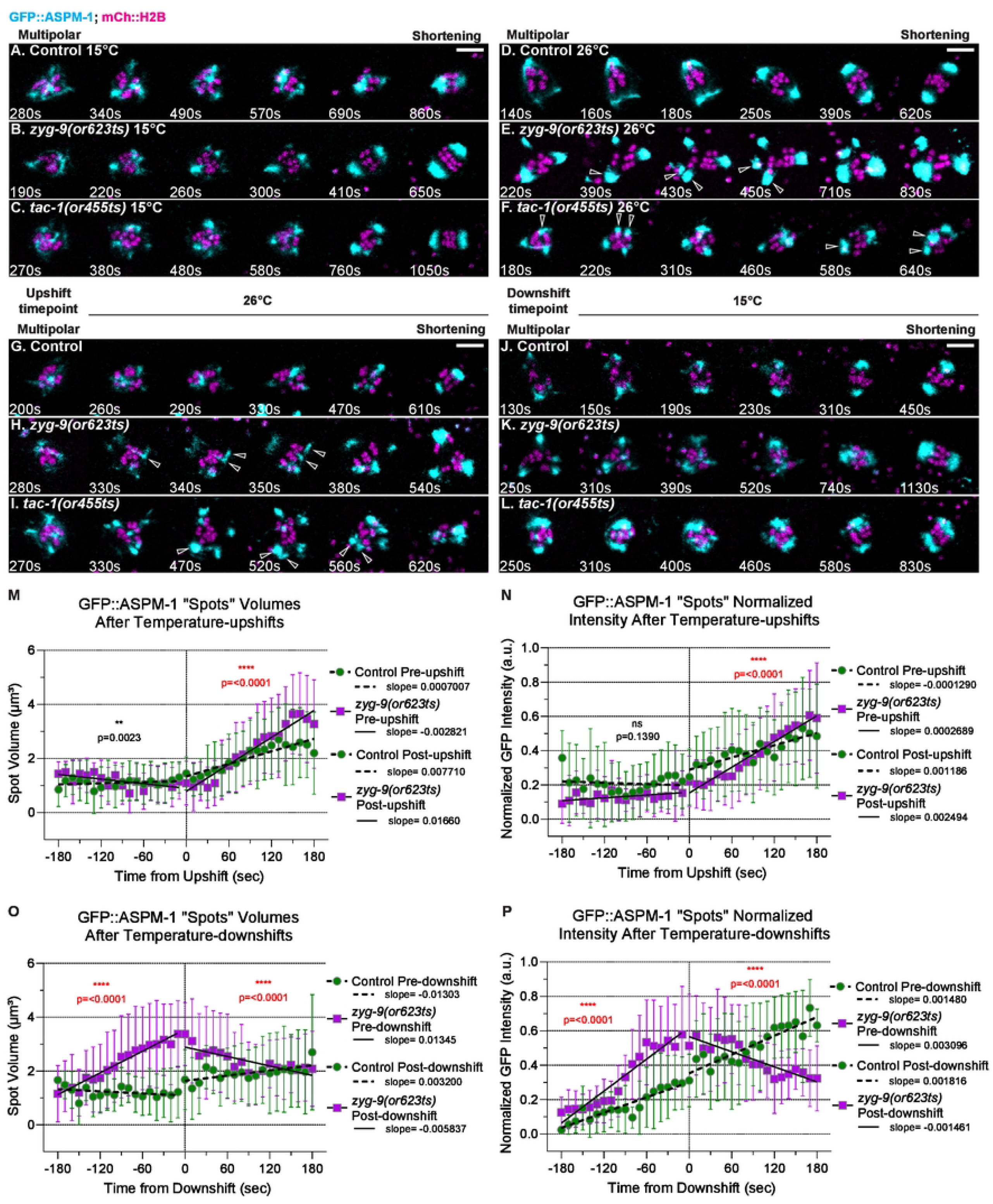
ZYG-9 and TAC-1 prevent the splitting of early pole foci and limit their growth during meiosis I bipolar spindle assembly. (A-L) Time-lapse maximum intensity projection images of live control and TS mutant oocytes expressing GFP::ASPM-1 and mCherry::H2B to mark spindle poles and chromosomes, at 15°C (A-C), at 26°C (D-F), and upshifted to 26°C (G-I) or downshifted to 15°C (J-L) during the multipolar stage. Montage frames highlight pole coalescence dynamics during the multipolar stage through to the onset of spindle shortening. (M-P) Quantification of control and *zyg-9(or623ts)* GFP::ASPM-1 foci volume and integrated pixel intensity (see Materials and Methods) pre- and post-multipolar upshift (M, N) and downshift (O,P). Slopes were compared using a two-tailed t-test to calculate P-values. **, P < 0.01; ****, P <0.0001. Scale bars = 5 μm.

To explore how ZYG-9 and TAC-1 influence pole coalescence, *zyg-9(or623ts)* and *tac-1(or455ts)* oocytes expressing the GFP::ASPM-1 pole marker were maintained at 15°C until the multipolar stage, and then upshifted to the restrictive temperature of 26°C, as described earlier (Materials and Methods). Preceding temperature upshifts in mutant oocytes, the early stages of pole foci coalescence appeared normal, with a diffuse network of small GFP::ASPM-1 foci surrounding the chromosomes. However, upon upshift the early pole foci appeared more dynamic in two ways. First, some pole foci split apart in all TS mutant oocytes after the upshifts, while no foci were observed to split apart after control upshifts (Fig. 4H, 4I, Table 1, Movie S10). Ultimately, mutant spindles frequently failed to become bipolar, in contrast to upshifted control oocytes (Fig. 2I, 4G). Second, pole foci grew more rapidly and to greater size after upshifts in *zyg-9(or623ts)* oocytes, compared to control oocytes (Fig. 4H, 4I, Fig. S5, Movies S11 & S12). Tracking GFP::ASPM-1 foci using Imaris revealed that while pole foci were similar in *zyg-9(or623ts)* and controls preceding upshift, after upshift the foci in *zyg-9(or623ts)* increased in both volume and intensity more rapidly than in control oocytes, indicated by a significant increase in the slopes of their growth rates and by the mutant foci ultimately becoming larger compared to controls (Fig. 4M, 4N). While we did not detect these differences in *tac-1(or455ts)* mutant oocytes (Fig. S6, Movie S13), the rapid growth of small GFP::ASPM-1 foci into more prominent foci after the temperature upshifts in *zyg-9(or623ts)* mutant oocytes suggests that ZYG-9 and TAC-1 may promote pole coalescence by limiting the growth of pole foci, in addition to preventing their splitting (see Discussion).

**Table 1.** ZYG-9 and TAC-1 prevent the splitting of early pole foci. Table of GFP::ASPM-1 splitting events in control and TS mutant oocytes upshifted and downshifted during the multipolar stage.

| GFP::ASPM-1 Splitting Events Post-upshift |  |
| --- | --- |
| Control | 0 of 6 oocytes (0,0,0,0,0,0) |
| *zyg-9(or623ts)* | 6, in 4 of 5 oocytes (1,2,2,1,0) |
| *tac-1(or455ts)* | 10, in 5 of 5 oocytes (1,1,2,2,4) |
| GFP::ASPM-1 Splitting Events up to 90 seconds Post-downshift |  |
| Control | 0 of 4 oocytes (0,0,0,0) |
| *zyg-9(or623ts)* | 6, in 5 of 5 oocytes (1,2,1,1,1)<sup>a</sup> |
| *tac-1(or455ts)* | 7, in 5 of 6 oocytes (0,1,2,2,1,1)<sup>b</sup> |
| GFP::ASPM-1 Splitting Events >90 seconds Post-downshift |  |
| Control | 0 of 4 oocytes (0,0,0,0) |
| *zyg-9(or623ts)* | 1, in 1 of 5 oocytes (0,0,1,0,0) |
| *tac-1(or455ts)* | 0 of 6 oocytes (0,0,0,0,0,0) |
| *Parentheses show the number of splitting events for each oocyte |  |
| <sup>a</sup> 3 of 6 splitting events occurred within 90 seconds after downshift |  |
| <sup>b</sup> 3 of 7 splitting events occurred within 90 seconds after downshift |  |

We also performed complimentary multipolar temperature-downshift experiments, as described above (Fig. S2), to assess the effects of restoring ZYG-9 and TAC-1 function to a disrupted early network of pole foci. Prior to the downshifts, we observed dynamic and prominent pole foci that grew larger and brighter in *zyg-9(or623ts)* oocytes compared to controls (Fig. 4K, 4L, Fig. S5, Movies S14-S16). Upon downshift, prominent pole foci reverted to a more diffuse network of smaller GFP::ASPM-1 foci, with GFP::ASPM-1 focus size and integrated pixel intensity decreasing following the downshift, showing an inverse effect compared to the upshifts, whereas poles in control oocytes showed a relatively constant growth trajectory (Fig. 4O, 4P). Also, nearly all examples of foci splitting in TS mutant oocytes occurred prior to or within 90 seconds of the downshifts (Table 1), indicating that restoring ZYG-9 or TAC-1 function rescues the stability of pole foci during coalescence. Finally, the downshifted poles coalesced into bipolar spindles that separated chromosomes into two equal sets in 4 of 5 *zyg-9(or623ts)*, and 6 of 6 *tac-1(or455ts)* oocytes (Table 1, Fig. S5). To summarize, these changes in pole dynamics after temperature upshifts and downshifts suggest that ZYG-9 and TAC-1 promote pole coalescence both by promoting pole stability and by limiting pole growth.

### ZYG-9 and TAC-1 suppress ectopic pole formation and maintain chromosome congression at the metaphase plate

We next asked whether ZYG-9 and TAC-1 are required for pole stability after a bipolar spindle has formed, using temperature upshifts during metaphase in TS mutant oocytes expressing either GFP::ASPM-1 or GFP::TBB-2 and mCherry::H2B. These temperature upshifts were done during an interval ranging from 8.7 to 18 minutes after cage onset and prior to spindle shortening/anaphase onset (Fig. S2). Following these temperature upshifts, the metaphase poles in TS mutant oocytes remained largely intact, but we nevertheless observed two defects. First, small ectopic poles appeared in the cytoplasm near the spindle in 5 of 17 *zyg-9(or623ts)*, 8 of 12 *zyg-9(or634ts)*, and 8 of 18 *tac-1(or455ts)* oocytes, while no such ectopic poles were observed in 19 control oocytes (Fig. 5A-D, Fig. 6A-D, Fig. S7 & S8, Movie S17). The ectopic poles in the mutant oocytes varied in size, were mobile and often fused with one of the two previously established poles before spindle shortening (Fig. 5L).

**FIGURE 5.**
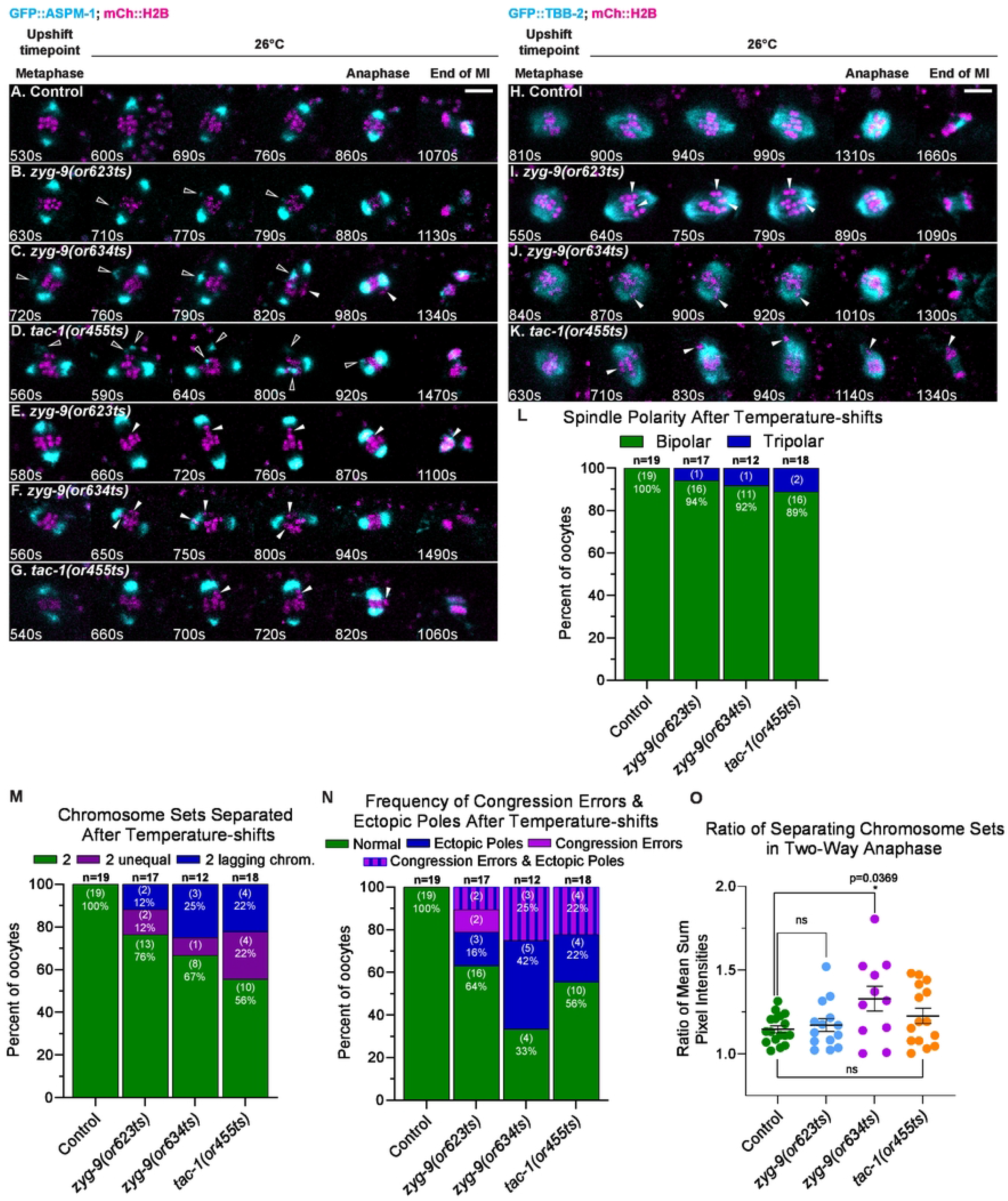
ZYG-9 and TAC-1 suppress ectopic spindle pole formation and maintain chromosome congression during meiosis I metaphase. (A-K) Time-lapse maximum intensity projection images of live control and TS mutant oocytes upshifted at metaphase and expressing either GFP::ASPM-1 and mCherry::H2B (A-G) or GFP::TBB-2 and mCherry::H2B (H-K). Montage frames highlight defects following metaphase upshift through to the end of meiosis I. White outlined arrowheads indicate ectopic spindle poles and solid white arrowheads indicate chromosome congression errors. (L-N) Number of spindle poles present at the onset of spindle shortening in metaphase upshifted oocytes (L); of chromosome congression errors (M); of ectopic poles in metaphase upshifted oocytes (M); of chromosome sets separated during anaphase (N) for metaphase upshifted oocytes; number of oocytes examined indicated above each bar, number scored with each phenotype shown in parentheses inside bars with corresponding percent below. (O) Ratio of fluorescence intensity of two separated chromosome sets in metaphase upshifted oocytes (see 5N) at the end of anaphase B. *, P <0.05. Scale bars = 5 μm.

**FIGURE 6.**
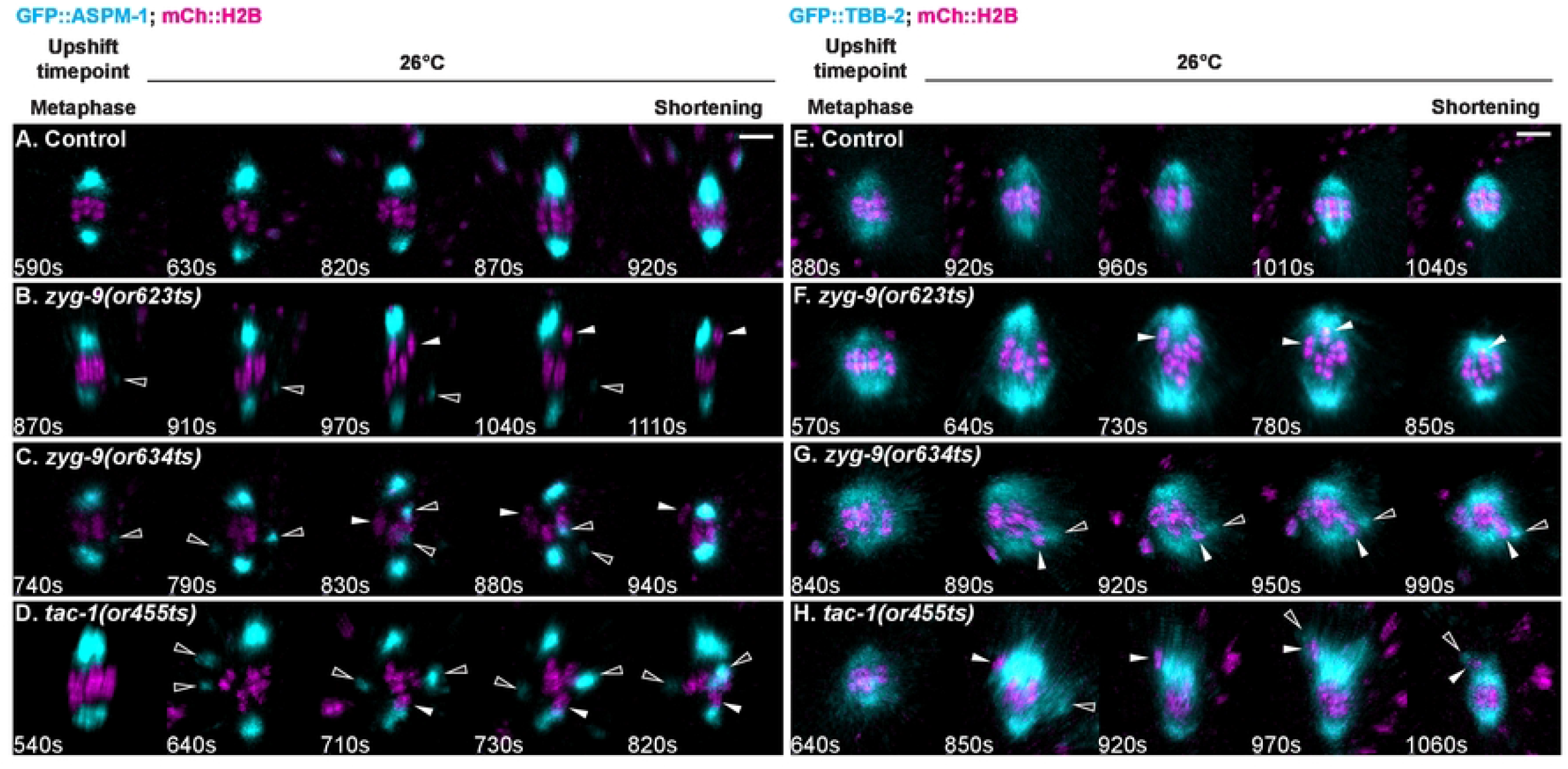
Ectopic poles are associated with chromosome congression defects in *zyg-9* and *tac-1* TS mutant oocytes. (A-H) Imaris rotated and snapshot projected time-lapse images (see Materials and Methods) of live control and TS mutant oocytes upshifted at metaphase expressing GFP::ASPM-1 and mCherry::H2B (A-D) or GFP::TBB-2 and mCherry::H2B (E-H). Montage frames highlight defects following metaphase upshift through to spindle shortening. White outlined arrowheads indicate ectopic spindle poles; solid white arrowheads indicate chromosome congression errors. Montages of oocytes in A, C, D, F, G, and H are the same oocytes shown in Figure 5 montages A, C, D, I, J, and K, respectively. Scale bars = 5 μm.

The second defect we observed in mutant oocytes after metaphase upshifts was a failure to maintain the congression of individual bivalents (paired homologs) at the metaphase plate. Individual bivalents moved away from the metaphase plate in 4 of 17 *zyg-9(or623ts)*, 3 or 12 *zyg-9(or634ts)*, and 4 of 18 *tac-1(or455ts)* metaphase-upshifted oocytes (Fig. 5E-G, 5I-K, Fig. 6B-H, Fig. S7 & S8, Movie S18). Importantly, the poorly congressed bivalents were associated with ectopic poles in 9 of the 11 TS mutant oocytes with congression errors (Fig. 5M, Fig. S8, Movie S19). Furthermore, in upshifted mutant oocytes expressing GFP::TBB-2 and mCherry::H2B and exhibiting congression defects, we observed ectopic microtubule bundles that extended toward poorly congressed bivalents, and spindle poles that appeared partially split (Fig. 6G-H, Fig. S8, Movie S20). These results are consistent with the hypothesis that either de novo ectopic pole formation, or pole instability, disrupts bipolar spindle structure and hence bivalent alignment. We therefore suggest that ZYG-9 and TAC-1 act during metaphase to suppress ectopic pole formation, and that this pole stability is important for maintaining chromosome congression.

We also asked whether the extensive anaphase chromosome separation defects observed in TS mutant oocytes kept at 26°C throughout meiosis I, and after RNAi knockdown (Chuang et al., 2020; Yang et al., 2003), are due to earlier pole coalescence and pole stability defects, or alternatively if ZYG-9 and TAC-1 might also have more direct roles in chromosome separation during anaphase. To distinguish between these two possibilities, we analyzed chromosome separation in TS mutant oocytes after metaphase upshifts and found that meiosis I chromosome separation outcomes resulting in just one or in three chromosome sets were nearly eliminated, suggesting that the earlier pole coalescence defects caused these more severe defects in chromosome separation (Fig. 5N). However, chromosome separation outcomes with two unequal chromosome sets were still observed in 2 of 17 *zyg-9(or623ts)*, 1 of 12 *zyg-9(or634ts)*, and 4 of 18 *tac-1(or455ts)* oocytes (Fig. 5N, 5O) and defects in the maintenance of chromosome congression accounted for all observed examples of chromosomes separating into two unequal sets (Movies S17 & S18). These results suggest that the earlier defect in pole coalescence and the later defect in pole stability together account for all defective chromosome separation outcomes. While we did not detect any further and possibly more direct roles for this protein complex during chromosome separation after metaphase upshifts, the modestly lower penetrance of the defects observed in TS mutant oocytes maintained at the restrictive temperature throughout meiosis I, compared to those observed after RNAi knockdown (see above), indicates that these TS alleles do not fully reduce gene function at the restrictive temperature. Thus we may not have been able to detect later requirements for lower levels of complex function.

### ZYG-9 and TAC-1 are required for anaphase spindle rotation and polar body extrusion

Previous studies of early embryonic mitosis have shown that loss of ZYG-9 or TAC-1 results in both short astral microtubules and abnormal mitotic spindle orientation due to the loss of uniform astral microtubule contact with the cell cortex (Bellanger and Gönczy, 2003; Bellanger et al., 2007). To determine if ZYG-9 and TAC-1 also influence meiotic spindle positioning, we scored spindle orientation relative to the overlying cell cortex in *zyg-9* and *tac-1* mutant oocytes after metaphase upshifts to the restrictive temperature. In control upshifted oocytes, the spindle shortened in the pole-to-pole axis and then rotated to become roughly perpendicular to the cortex, such that three or more of the six homologous chromosome pairs contacted the cortex prior to chromosome separation (Fig. 7A, 7E; 19/19 oocytes), one measure of proper spindle rotation (Yang et al., 2003, 2005; Crowder et al., 2015; Vargas et al., 2019). In contrast, spindles failed to rotate, or only partially rotated, in 4 of 13 *zyg-9(or623ts)*, 8 of 12 *zyg-9(or634ts)*, and 8 of 18 *tac-1(or455ts)* oocytes (Fig. 7B-E, Fig. S9, Movie S21). We also scored spindle orientation by measuring the angle of the spindle axis relative to a tangent of the cortex after rotation (Vargas et al., 2019). While all control oocytes rotated to within an 80-90° range, most of the spindles in *zyg-9* and *tac-1* mutant oocytes failed to fully rotate (Fig. 7F).

**FIGURE 7.**
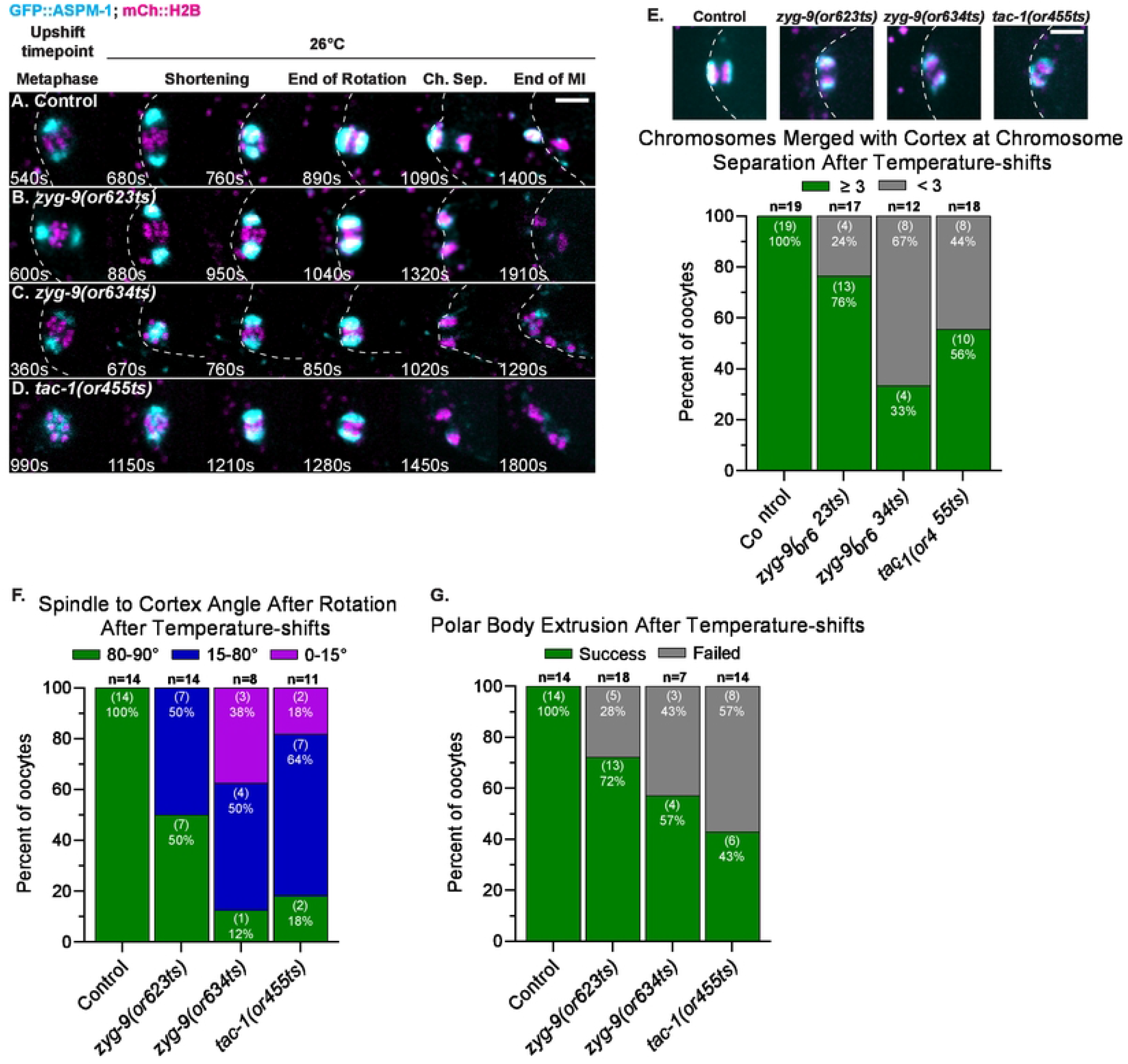
ZYG-9 and TAC-1 are required for meiosis I anaphase spindle rotation and polar body extrusion. (A-D) Time-lapse maximum intensity projection images of live control and TS mutant oocytes expressing GFP::ASPM-1 and mCherry::H2B upshifted at meiosis I metaphase; polar body extrusion failed in all three mutant oocytes. Oocytes were rotated so that the cortex was positioned to the left of the spindle; dashed lines depict the oocyte cortex. (E) Early anaphase time-lapse images of live control and TS mutant oocytes after metaphase upshifts. Bar graph quantifies spindle rotation based on the number of chromosomes adjacent to the cortex at the start of chromosome separation in metaphase upshifted oocytes (see Materials and Methods). (F) Spindle angles relative to the cortex at early anaphase in metaphase upshifted oocytes. (G) Number of control and TS mutant metaphase upshifted oocytes that extrude a polar body during meiosis I (see Materials and Methods). Scale bars = 5 μm.

Because spindle rotation is thought to facilitate extrusion of excess oocyte chromosomes into a polar body, we also scored whether chromosomes were successfully extruded. In control upshifted oocytes, chromosomes were always detected within an external polar body at the onset of meiosis II (n = 14). By contrast, polar body extrusion was defective after metaphase upshifts in 5 of 18 *zyg-9(or623ts)*, 3 of 7 *zyg-9(or634ts)*, and 8 of 14 *tac-1(or455ts)* oocytes, with all chromosomes present within the oocyte cytoplasm at the beginning of meiosis II (Fig. 7B-E, 7G, Fig. S9, Movie S22). While polar body extrusion was more likely to fail in oocytes with spindle rotation defects, the correlation was only partial (Fig. S9I), suggesting that other defects may contribute to the extrusion failures.

### ZYG-9 and TAC-1 also promote pole coalescence during meiosis II spindle assembly

Many of the proteins required for oocyte meiosis I are present during and thought to be required for meiosis II, although evidence for this is lacking as meiosis I requirements can preclude the assessment of later requirements. We therefore used our fast-acting TS alleles to ask whether ZYG-9 and TAC-1 are required for meiosis II spindle assembly. The second meiotic division begins immediately after extrusion of the first polar body and spindle assembly largely resembles meiosis I except that (i) the spindle is smaller and the duration of meiosis II shorter (McNally and McNally, 2005), and (ii) the nuclear envelope has already disassembled, with microtubules appearing amongst the chromosomes instead of forming a peripheral cage structure (Wolff et al., 2016). To view normal meiosis II pole assembly dynamics, we imaged control and TS mutant oocytes from strains expressing GFP::ASPM-1 and mCherry::H2B and maintained at 15°C. Meiosis II began with small GFP::ASPM-1 pole foci forming around the chromosomes and then coalescing into a bipolar spindle with pairs of sister chromatids aligned between the poles (Fig. 8A-D, Fig. S10). We next upshifted control and TS mutant oocytes to the restrictive temperature during meiosis I polar body extrusion, to inactivate ZYG-9 and TAC-1 throughout all of meiosis II while bypassing the meiosis I requirements. Spindle pole foci in *zyg-9* and *tac-1* upshifted oocytes also appeared around the chromosomes but were more dynamic and took longer or failed to assemble into bipolar spindles (Fig. 8F-H, Fig. S10). While the spindle bipolarity and chromosome separation defects in all three mutants were reduced in penetrance during meiosis II compared to meiosis I (Fig. 1I, 1J; Fig. 8I, 8J), these results indicate that ZYG-9 and TAC-1 also influence pole coalescence and spindle bipolarity during meiosis II.

**FIGURE 8.**
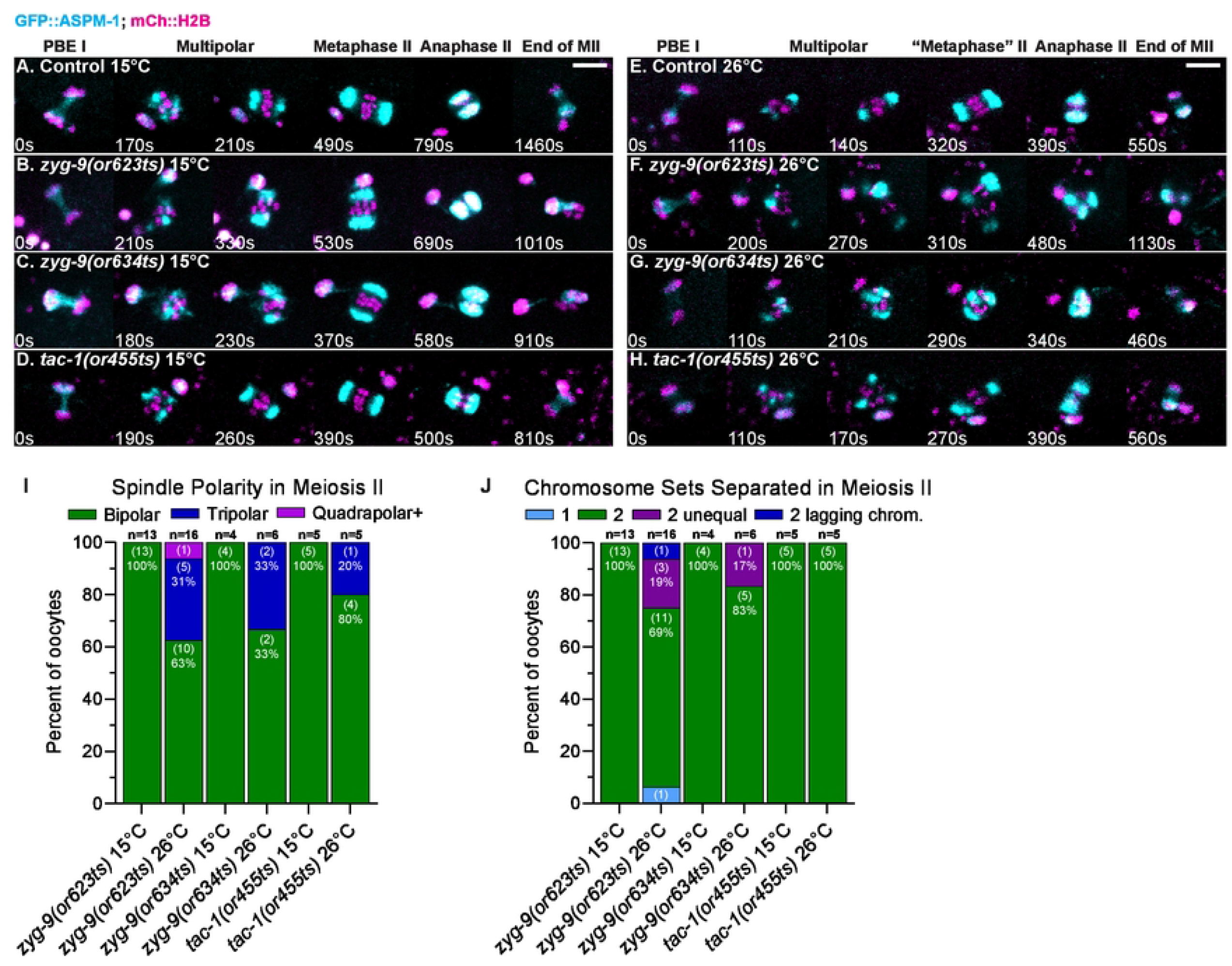
ZYG-9 and TAC-1 are required for meiosis II spindle assembly. (A-H) Time-lapse maximum intensity projection images during meiosis II of live control and TS mutant oocytes expressing GFP::ASPM-1 and mCherry::H2B at 15°C (A-D) and at 26°C (E-H). t=0 is the timepoint when meiosis I chromosome separation ends. (I-J) Number of spindle poles present per oocyte at the onset of spindle shortening (I); of chromosome sets separated during anaphase II (J); number of oocytes examined indicated above each bar, number scored with each phenotype shown in parentheses inside bars with corresponding percent below. Scale bars = 5 μm.

## Discussion

Gene requirements for the acentrosomal assembly of oocyte meiotic spindles have been investigated to varying extents in different model organisms (Dumont and Desai, 2012; Severson et al., 2016; Mullen et al., 2019; Ohkura, 2015). Experiments in *Drosophila, C. elegans* and vertebrate oocytes have shown that XMAP215 orthologs, and their TACC-domain containing binding partners, have important roles in acentrosomal oocyte spindle assembly.

Non-conditional RNAi knockdowns in *C. elegans* have shown that XMAP215/ZYG-9 and TACC/TAC-1 are required for multiple aspects of meiosis I spindle assembly: preventing microtubule bundles from crossing through the space occupied by oocyte chromosomes after nuclear envelope breakdown, limiting the overall levels of spindle- and cortex-associated microtubules, and promoting the coalescence of early pole foci into a bipolar structure that separates chromosomes into two equal sets. Our results, using fast-acting TS alleles of *zyg-9* and *tac-1* with live imaging and fluorescent protein fusions, indicate that the later pole coalescence defects in *zyg-9* and *tac-1* mutants occur independently of the earlier defect in microtubule cage structure, and that ZYG-9 and TAC-1 also are required for pole coalescence during meiosis II. Our results further suggest that ZYG-9 and TAC-1 act during metaphase to promote acentrosomal pole stability, and that this metaphase pole stability contributes to the maintenance of chromosome congression at the metaphase plate during oocyte meiotic cell division. Another recent study also reports data indicating that ZYG-9 and TAC-1 are required for pole stability during oocyte meiosis I, and in addition shows that ZYG-9 and TAC-1 are codependent each other for their localization to oocyte meiotic spindle poles (Mullen et al., 2022).

### ZYG-9 and TAC-1 have multiple and temporally distinct requirements during oocyte meiotic spindle assembly

Our use of fast-acting TS alleles has identified at least three temporally separable requirements for ZYG-9 and TAC-1 during oocyte meiotic cell division. First, ZYG-9 and TAC-1 limit the appearance of microtubule bundles to the periphery after nuclear envelope breakdown such that they form a roughly spherical cage that surrounds the oocyte chromosomes early in spindle assembly. Second, our temperature upshift and downshift experiments indicate that the later defect in pole coalescence does not depend on an earlier cage stage defect. Following temperature upshifts during prometaphase pole coalescence, mutant oocytes exhibited polarity defects even in the absence of any pre-existing cage defect. Furthermore, following temperature downshifts at the prometaphase stage, mutant spindles with early cage defects formed bipolar spindles that separated chromosomes normally into two equally sized sets, indicating that a pre-existing cage defect is not sufficient to cause later defects in pole coalescence. We conclude that the defective coalescence of pole foci in TS mutant oocytes represents a distinct and later requirement for this protein complex during the coalescence process itself. Our results further indicate that the mutant spindle polarity defects are not due to coalescence occurring in three dimensions, rather than within the more restricted two-dimensional surface normally defined by the microtubule bundles that form the early cage structure, as we previously speculated (Chuang et al., 2020). Finally, following temperature upshifts after the completion of meiosis I, we also observed pole coalescence defects during meiosis II, a third temporally distinct requirement. We conclude that this protein complex is active throughout meiosis I and II and performs multiple temporally separable and therefore more direct roles.

We also have shown that the pole stability defects in *zyg-9* and *tac-1* mutant oocytes appear to be independent of the overall increase in spindle-associated microtubule levels observed after ZYG-9 or TAC-1 RNAi knockdowns (Chuang et al., 2020). While we documented similar pole coalescence and chromosome separation defects at the restrictive temperature in all three TS alleles, spindle-associated microtubule levels were reproducibly elevated only in *tac-1(or455ts)* oocytes, indicating that the pole coalescence defects do not depend on an overall increase in microtubule levels. However, these results do not rule out more localized or temporally restricted roles for destabilizing microtubules during pole coalescence and spindle assembly (see below). Finally, the requirements for ZYG-9 and TAC-1 to limit oocyte spindle and cortex microtubule levels during meiosis I further indicates that this protein complex likely functions throughout much of oocyte meiotic cell division to influence microtubule stability and cell division.

### An emerging role for the regulation of microtubule dynamics during polar body extrusion

We have shown that both rotation of the oocyte spindle, which normally makes the spindle axis perpendicular to the overlying cell cortex, and extrusion of chromosomes into a polar body, often fail in *zyg-9* and *tac-1* mutant oocytes. These defects were observed after metaphase upshifts and thus appear to be independent of the earlier cage structure and prometaphase pole coalescence defects. *C. elegans* oocyte meiotic spindle rotation is thought to require sparse and short astral microtubules, microtubule motors, and shortening of the spindle into a near-spherical shape (Vargas et al., 2019; Crowder et al., 2015; Fabritius et al., 2011). Defects in spindle structure or dynamics that occurred after the metaphase upshifts and prior to rotation and extrusion, including the transient appearance of ectopic pole foci and misshapen spindle poles, might indirectly account for the rotation and extrusion defects.

While spindle rotation defects might also indirectly cause failures in polar body extrusion, contractile ring assembly and ingression have not been examined in mutants with oocyte spindle rotation defects, and whether rotation is required for proper extrusion is not known (Schlientz and Bowerman, 2020). Furthermore, the lack of a clear correlation between failed rotation and failed extrusion after metaphase upshifts in TS mutant oocytes, with polar body extrusion sometimes failing after apparently normal rotation, suggest that ZYG-9 and TAC-1 could have additional and more direct roles in polar body extrusion. Recent studies of the *C. elegans* TOG domain containing protein and CLASP2 family member CLS-2 indicate that proper regulation of microtubule stability is important for contractile ring assembly and ingression during polar body extrusion (Schlientz and Bowerman, 2020). However, in contrast to *zyg-9* and *tac-1* mutant oocytes, in which strong loss of function results in abnormally high levels of spindle- and cortex-associated microtubules during oocyte meiotic cell division, loss of CLS-2 results in reduced microtubule levels. It will be interesting to further explore the requirements for ZYG-9 and TAC-1 during spindle rotation and polar body extrusion, with later temperature upshifts and downshifts, and live imaging of contractile ring assembly and ingression.

### ZYG-9 and TAC-1 prevent the splitting and limit the growth of pole foci during prometaphase pole coalescence

Our temperature upshift and downshift experiments indicate that ZYG-9 and TAC-1 act during prometaphase to promote the coalescence of pole foci into a bipolar spindle.

Temperature upshifts during pole coalescence, with live imaging and the GFP::ASPM-1 spindle pole marker, resulted in two informative changes in pole dynamics. First we often saw early GFP::ASPM-1 pole foci split in two after pole coalescence upshifts in TS mutant oocytes, while we did not observe such splitting after control oocyte upshifts. We conclude that ZYG-9 and TAC-1 promote the fusion of early pole foci in part by opposing their fission, with fusion over time reducing their number and resulting in bipolarity. We were surprised to also observe a second change in *zyg-9(or623ts)* oocytes: a rapid increase in both the average size and the integrated pixel intensity of the GFP::ASPM-1 foci after upshifts during pole coalescence, and a corresponding decrease in their size and intensity after pole coalescence downshifts. While we observed these effects on pole dynamics only after temperature shifts in *zyg-9(or623ts)* and not in *tac-1(or455ts)* oocytes, we did observe the splitting of pole foci after *tac-1(or455ts)* upshifts. We speculate that these conditional and partial loss-of-function alleles differently alter complex function, and the failure to detect the changes in pole foci growth properties in a partial loss-of-function *tac-1* mutant does not preclude such a role for the ZYG-9/TAC-1 complex. We conclude that proper pole coalescence depends not only on the stability of coalescing pole foci but also on their growth dynamics.

### ZYG-9 and TAC-1 promote the maintenance of spindle bipolarity and chromosome congression during metaphase

We observed two additional and correlated phenotypes upon later metaphase upshifts in TS mutant oocytes, after bipolar oocyte spindles had already assembled. Ectopic pole foci often appeared near pre-existing bipolar spindles, and individual pairs of homologous chromosomes often migrated away from the metaphase plate toward one pole. While we did not observe any splitting of spindle poles to produce these ectopic foci, we cannot rule out such fission as their source, as opposed to de novo assembly. Notably, these ectopic foci were almost always observed in association with homologous chromosome pairs, or bivalents, that moved away from the metaphase plate: in 9 of 11 oocytes with bivalents that became displaced towards one pole after the metaphase upshifts, we also observed an ectopic focus of GFP::ASPM-1 or an ectopic bundle of spindle microtubules. We speculate that similar such defects escaped detection, due to the limits of our imaging methods, but were also associated with displaced bivalents in the remaining two oocytes. We conclude that this protein complex promotes both the prometaphase coalescence of pole foci and the subsequent stability of spindle poles during metaphase.

### Microtubule elongation and acentrosomal spindle pole coalescence

Presumably the prometaphase pole coalescence and metaphase pole stability defects in *zyg-9* and *tac-1* mutant oocytes reflect a requirement for this protein complex in the regulation of microtubule stability (see Introduction). Most studies of XMAP215 and TACC orthologs have focused on their roles in promoting microtubule stability, often acting as microtubule polymerases (Akhmanova and Steinmetz, 2015; Gunzelmann et al., 2018; Thawani et al., 2018; Cook et al., 2019). However, ZYG-9 and TAC-1 appear at least in sum to promote microtubule instability during oocyte meiotic cell division (Chuang et al., 2020), with loss of their function resulting in elevated oocyte microtubule levels, both in association with the meiotic spindle and throughout the oocyte cortex. This is in substantial contrast to most studies of XMAP215 and TACC orthologs, and to ZYG-9 and TAC-1 during mitosis in the early *C. elegans* embryo, when loss of their function results in decreased microtubule levels (see Introduction).

While our results indicate that the pole coalescence and stability defects appear to be independent of an overall increase in microtubule levels during oocyte meiotic cell division (see above), we nevertheless speculate that spatially and temporally restricted limits to microtubule elongation might underlie the defects in pole coalescence and stability observed in mutant oocytes. For example, XMAP215 orthologs are enriched at meiotic and mitotic spindle poles, where they have been proposed to both stabilize microtubule minus ends and counter-act the activity of microtubule depolymerases at plus ends (Lee et al., 2001; Peset et al., 2005; Tournebize et al., 2000). If ZYG-9 and TAC-1 can also act at pole foci, but to destabilize minus ends or limit microtubule polymerization at plus ends, localized and excessive microtubule growth in mutant oocytes might destabilize coalescing pole foci, with the forces generated by microtubule elongation pushing foci apart. Such spatially and temporally localized perturbations to microtubule dynamics in the TS mutant oocytes might not be reflected in overall microtubule levels. Similarly, excessive microtubule polymerization from spindle poles during metaphase might also promote pole instability and thereby generate ectopic pole foci that mislocalize congressed bivalents. It is perhaps less clear how limiting the rate of pole foci growth might promote coalescence, but excessively rapid foci growth might similarly promote locally excessive microtubule polymerization in mutant oocytes.

### Liquid-liquid phase transitions and acentrosomal spindle pole coalescence

ZYG-9 and TAC-1 orthologs can undergo liquid-liquid phase transitions, and such condensate formations might also mechanistically contribute to acentrosomal pole coalescence and stability in *C. elegans* oocytes. Recent studies in non-human mammalian oocytes have shown that XMAP215/chTOG and TACC/TACC3 exhibit liquid droplet properties during oocyte meiotic spindle assembly (So et al., 2019). Intriguingly, *C. elegans* XMAP215/ZYG-9, in association with *in vitro* condensates of the coiled-coil centrosome scaffolding protein SPD-5, can concentrate tubulin and promote microtubule nucleation in the absence of any other factors (Woodruff et al., 2017). However, *C. elegans* ZYG-9 and TAC-1 are generally enriched at oocyte meiotic spindle poles and also broadly associated with spindle microtubules (Chuang et al., 2020), but do not appear to exhibit any of the condensate dynamics observed for chTOG and TACC3 in non-human mammalian oocytes; such condensates also are not observed in human oocytes (So et al., 2019).

Liquid droplet properties of spindle proteins during oocyte meiotic spindle assembly have not been documented in *C. elegans*, but pole foci coalescence with the GFP::ASPM-1 marker does somewhat resemble the fusion of condensate droplets reported in mammalian oocytes. Notably, depletion of TACC3—but not chTOG—from mouse oocytes disrupts the ability of other spindle proteins to form condensates, with liquid-liquid phase transition appearing to play an important role in mammalian oocyte spindle pole assembly. Perhaps the absence of ZYG-9 or TAC-1, even if they do not obviously appear to form condensates themselves during *C. elegans* oocyte spindle assembly, nevertheless can alter the properties of condensates that are involved in pole coalescence, such that they are less stable and more prone to splitting or nascent assembly, and to rapid growth. Higher spatial and temporal resolution imaging of ZYG-9 and TAC-1, and other factors, during oocyte meiotic spindle assembly may provide further insight.

### ZYG-9 and TAC-1 promote microtubule instability during oocyte meiotic cell division

The negative regulation of overall microtubule levels by ZYG-9 and TAC-1 in *C. elegans* oocytes is in stark contrast to its subsequent role in promoting microtubule stability during the first embryonic mitosis, which immediately follows the completion of oocyte meiosis I and II. Furthermore, most studies in other model systems, both in vivo and in vitro, have focused on XMAP215 and TACC roles as microtubule stabilizing factors (Akhmanova and Steinmetz, 2015). For example, budding yeast XMAP215/Stu2p has been shown to synergize with gamma-tubulin to promote microtubule nucleation and polymerization (Gunzelmann et al., 2018; Thawani et al., 2018), and XMAP215 orthologs are often described as microtubule polymerases (Akhmanova and Steinmetz, 2015; Cook et al., 2019; Brouhard et al., 2008). While it is possible that ZYG-9 and TAC-1 also promote microtubule stability during oocyte meiotic cell division, the substantial increase in both spindle and cortex-associated microtubule levels after strong reduction of either ZYG-9 or TAC-1 clearly indicates a predominantly negative role during oocyte meiotic cell division.

Though less extensively studied, some XMAP215 family members also destabilize microtubules in some contexts. Studies of budding yeast XMAP215/Stu2p both in vitro and in vivo indicate that it can limit microtubule length and decrease the rate of microtubule catastrophe (van Breugel et al., 2003; Kosco et al., 2001), while studies of interphase microtubules in *Drosophila* S2 cells suggest that XMAP215/Mini-spindles negatively regulates pauses in microtubule growth, although this role may be independent of D-TACC (Brittle and Ohkura, 2005). Finally, in vitro studies using purified proteins or *Xenopus* extracts have shown that XMAP215 can promote both microtubule stability (Tournebize et al., 2000; Vasquez et al., 1994; Brouhard et al., 2008) and microtubule instability (Shirasu-Hiza et al., 2003). *C. elegans* oocyte meiotic spindle assembly provides a particularly appealing model system for further investigating the role of XMAP215 and TACC family members as negative regulators of microtubule stability, and for how they can transition from being negative to positive regulators. Lastly, whether XMAP215 and TACC family members act exclusively positively or negatively to regulate microtubule stability in any one setting, or perhaps instead use a balance of positive and negative regulation in some contexts, requires further investigation.

### Conservation of oocyte spindle assembly mechanisms across animal phyla

Studies of XMAP215 and TACC orthologs during meiotic cell division in *Drosophila* and mammalian oocytes indicate that their roles during acentrosomal spindle assembly may not be fully conserved, as the defects in some cases resemble those we have observed in *C. elegans*, and in other cases appear distinct. In *Drosophila* oocytes, XMAP215/Mini-spindles and D-TACC also are enriched at spindle poles, and loss of either results in many oocytes assembling tripolar spindles that are remarkably similar to the tripolar spindles we frequently observed in *zyg-9* and *tac-1* mutants. However, Mini-spindles and D-TACC require the minus-end directed kinesin 14 family member Ncd for their localization to spindle poles (Cullen and Ohkura, 2001), and Ncd mutant oocytes also often assemble tripolar spindles (Sköld et al., 2005). The localization of ZYG-9 and TAC-1 in *klp-15/16* mutant oocytes has not been reported, but loss of the kinesin 14 family members KLP-15/16 in *C. elegans* results in mutant oocytes with a pole coalescence phenotype that differs substantially from the *zyg-9* and *tac-1* mutant phenotype (Chuang et al., 2020), suggesting that ZYG-9 and TAC-1 may not function so directly with kinesin 14 family members as do their *Drosophila* orthologs.

In an extensive survey of mammalian oocyte spindle proteins, XMAP215/chTOG and TACC3 and several other spindle pole proteins were shown to undergo phase transitions and be present in spindle-associated condensates (So et al., 2019). However, as noted earlier, GFP fusions to ZYG-9 and TAC-1 do not appear to form condensates *in C*. elegans oocytes but are more diffusely localized throughout the spindle with some enrichment at the poles (Chuang et al., 2020). Moreover, in contrast to the elevated levels of microtubules and the extensive defects in spindle bipolarity observed in *zyg-9* and *tac-1* mutant oocytes, bipolar spindles assembled with significantly reduced spindle microtubule levels after depletion of chTOG or TACC3 from mouse oocytes, and no defects in pole assembly or spindle bipolarity were reported (So et al., 2019). Nevertheless, it is intriguing that in mouse oocytes depleted of endogenous TACC3, but expressing a TACC3 construct lacking the N-terminal residues required for phase separation, oocyte meiotic poles were destabilized (So et al., 2019). Finally, normal human oocytes are exceptional in exhibiting dramatic pole instability and chromosome separation defects relative to other mammalian oocytes, due at least in part to loss of expression of KIFC1/kinesin 14 (So et al., 2022). In *C. elegans*, loss of KLP-15/16/kinesin 14 does result in failed pole coalescence but does not result in pole stability defects like those observed in *zyg-9* and *tac-1* mutant oocytes (Chuang et al., 2020), and in normal human oocytes (So et al., 2022). While there appear to be both substantial similarities and differences in the requirements for oocyte meiotic spindle assembly in different species, higher spatial and temporal resolution studies of pole coalescence and spindle assembly dynamics in live control and mutant oocytes will likely reveal further conservation and divergence of mechanism.

### Temporal dissection of gene requirements

Fast-acting, temperature-sensitive alleles provide a uniquely powerful tool for high-resolution temporal dissection of gene requirements and are particularly useful for investigating the relatively rapid oocyte meiotic and early embryo mitotic cell divisions in *C. elegans*. Coupled with devices that can control and rapidly change sample temperature during live imaging, fast-acting TS alleles have provided insights into both mitotic (Severson et al., 2000; Davies et al., 2014; Jordan et al., 2016) and meiotic (McNally et al., 2014, 2016; Connolly et al., 2014, 2015) cell division in *C. elegans*. However, while large-scale screens have identified thousands of temperature-sensitive, embryonic-lethal *C. elegans* mutants, relatively few have defects in early embryonic cell division (Pintard and Bowerman, 2019). Furthermore, temperature-sensitive alleles have been identified in only a few hundred of the roughly 2000 *C. elegans* genes known to be essential (Pintard and Bowerman, 2019), and only about half of the alleles in a collection of TS mutants with early embryonic cell division defects were found to be fast-acting (O’Rourke et al., 2011). While still valuable, slower-acting TS alleles require turnover of the encoded proteins and are therefore generally not useful for temporally dissecting gene requirements during the rapid early embryonic cell divisions in *C. elegans*.

The recent development of CRISPR-mediated degron tagging of endogenous loci for temporally controlled, auxin-mediated protein degradation now provides a powerful tool for engineering conditional loss of function alleles throughout the *C. elegans* genome (Zhang et al., 2015; Dickinson and Goldstein, 2016). Degron tagging also provides impressive temporal resolution, with tagged proteins substantially knocked down after only 20 to 30 minutes of exposure to auxin (Zhang et al., 2015; Divekar et al., 2021). However, fast-acting TS alleles can reduce function much more rapidly, within seconds or minutes. With further advances in the increasingly powerful ability to predict protein structure based on primary amino acid sequence, perhaps it will become feasible to accurately predict missense mutations that are likely to generate fast-acting, temperature-sensitive mutations, adding to our ability to temporally manipulate gene function in model organisms.

## Materials and Methods

### *C. elegans* strain maintenance

*C. elegans* strains used in this study are listed in Table S1. Temperature-sensitive and control strains were constructed and maintained at 15°C using previously described culturing methods (Brenner, 1974).

### Image acquisition

All imaging was performed using a Leica DMi8 microscope outfitted with a spinning disk confocal unit–CSU-W1 (Yokogawa) with Borealis (Andor), dual iXon Ultra 897 (Andor) cameras, and a 100x HCX PL APO 1.4–0.70NA oil objective lens (Leica). Metamorph (Molecular Devices) imaging software was used for controlling image acquisition. The 488nm and 561nm channels were imaged simultaneously every 10s with 1μm Z-spacing for a total stack size of 20μm.

*In utero* live imaging of oocytes was accomplished by mounting young adult worms with single row of embryos to a 5% agarose pad on a 24×40mm glass coverslip with 1.5 µl each of 15°C M9 buffer and 0.1 µm polystyrene microspheres (Polysciences Inc.), and were gently covered with a 18×18mm coverslip. Samples were then transferred to a fluidic temperature-controlled CherryTemp chip (CherryBiotech) that enabled imaging at precisely the indicated temperature and rapid temperature-shift experiments.

### Temperature-shift experiments

For multipolar upshift and downshift experiments, temperature-shifts were performed after ovulation and preceding spindle bipolarity and fell within a range of 2.7 to 6.8 minutes after cage onset. For metaphase upshift experiments, temperature-shifts were performed after spindles achieved bipolarity and aligned chromosomes and preceding spindle shortening and fell into a range of 8.7 to 18 minutes after cage onset. Upshifted oocytes that did not meet these criteria or that subsequently arrested were excluded from analysis. When analyzing the time intervals for multipolar and metaphase upshifts relative to spindle shortening, both control and TS mutant oocytes at 26°C progressed through meiosis I much faster than at 15°C, as expected (Fig. S11). Similarly, oocytes upshifted at the multipolar stage reached anaphase onset more quickly than oocytes upshifted at metaphase (Fig. S11). To describe our temperature upshift and downshift experiments in all figures, we define t=0 as the first frame prior to the appearance of a cage structure.

### Image processing and analysis

General image processing including merging and cropping of red/green channels and z-projected images was done through FIJI (Schindelin et al., 2012). All time lapse images shown in the figures are Maximum Intensity Projections of all focal planes unless otherwise noted. To account for differences in movie signal quality inherent to *in utero* live cell imaging, the intensity scales for montages in all figures were set individually to give the clearest depiction of the spindle and chromosomes; all pixel intensity based quantification was done using the raw unadjusted pixel values. The time lapse images in Figure 6 and S8 were obtained after rotation of 3-dimensional images using Imaris, with Imaris snapshots (roughly equivalent to Maximum Intensity Projections) used to acquire images of the rotated stacks. The montage frames for each stage were chosen as follows: “NEBD” is the timepoint immediately preceding the appearance of microtubule bundles forming the cage structure and serves as t=0; “Cage” is the timepoint that best displays the spherical cage-like network of microtubules that forms around the chromosomes; “Multipolar” is the timepoint that best displays the coalescing microtubule network after the cage stage and ovulation but preceding spindle bipolarity; “Metaphase” is the timepoint that best displays the bipolar spindle with chromosomes aligned at the metaphase plate and preceding spindle shortening; as metaphase is unidentifiable in TS mutant spindles at 26°C, the “Metaphase” frames in those montages are instead from the timepoint best displaying abnormal spindle morphology before the start of spindle shortening; “Anaphase” is a timepoint during anaphase A showing the shortened spindle and preceding chromosome separation; “End of MI” is a timepoint after chromosome separation that best displays the number of chromosome sets separated during anaphase. For montages of temperature-shift experiments, the frame at the point of upshift/downshift is always shown. For montages of meiosis II, frames for each stage were chosen as in meiosis I except “PBE I” is the timepoint where meiosis I chromosome separation ends and serves as t=0.

To quantify the ratio of mCherry::H2B signal of chromosome sets in two-way anaphases, a sum projection of the red channel was created in FIJI and then background subtracted using a 50 pixel rolling ball radius. A region of interest was drawn around each chromosome mass at the end of chromosome separation and a ratio was made of the mean sum intensity.

Normalized microtubule pixel intensity was quantified in FIJI as described previously (Chuang et al., 2020). For normalized microtubule intensity quantification by meiotic stage (Fig. 3A), measurements from three consecutive time points were taken at each stage per oocyte and reported as an average. The stages are defined as follows: the “Cage” is when GFP::TBB-2 marked microtubules formed a spherical cage-like network around the chromosomes; “Multipolar” is the timepoint midway between the Cage and Metaphase stages; “Metaphase” is 30 seconds before the onset of spindle shortening; “Anaphase” is the halfway point from spindle shortening onset to the start of chromosome separation; “Telophase” is midway through chromosome separation. For combined normalized microtubule intensities (Fig. 3B), the microtubule intensity values for all meiotic stages as quantified for Fig. 3A were taken from each oocyte and grouped by genotype.

Three-dimensional projection and rotation movies were made using Imaris software (Bitplane) and were used to analyze cage structures (Fig. 2K), assess pole splitting (Table 1), score spindle pole numbers (Figs. 1I, 2I, 5L, 8I), and analyze chromosome separation outcomes (Figs. 1J, 2J, 5N, 8J) and polar body extrusion (Fig. 7G).

A cage structure was scored as defective when GFP::TBB-2 marked microtubule bundles passed through the interior of the nucleus instead of maintaining a roughly spherical cage around the chromosomes (Fig. 2K; Movies S7 & S9).

For assessing pole splitting (Table 1), GFP::ASPM-1 marked foci were monitored from coalescence until spindle shortening and a splitting event was defined as when a single focus fissions into two or more separate foci (Movie S10).

To quantify spindle pole numbers at the time of spindle shortening, we counted the number of major GFP::TBB-2 microtubule asters or GFP::ASPM-1 pole foci (Figs. 1I, 2I, 5L, 8I).

To evaluate anaphase outcomes (Figs. 1J, 2J, 5N, 8J), we recorded the number of mCherry::H2B marked chromosome sets in each oocyte at the end of meiosis I or II.

To quantify GFP::ASPM-1 foci volumes and intensities (Fig. 4M-P), we used the Spots creation tool in Imaris (Movie S23). In the Spots creation wizard, we enabled “different spot sizes (region growing)” to capture foci across a range of sizes. The source channel for detection was set to “Green Channel” and the estimated XY diameter for Spot detection was 1.47μm. The background subtraction (local contrast) method was used to threshold foci, and the threshold value was adjusted manually until Spots appeared over the GFP::ASPM-1 foci of the forming oocyte spindle. During Spot detection, the region growing diameter was measured from the region volume. To eliminate off-targets, we filtered the Spot creation to exclude Spots further than 10μm from the chromosomes marked by mCherry::H2B and then manually deleted remaining off-target Spots. Spot volume and intensity data from each movie were then exported to Microsoft Excel (Microsoft), the intensity data were normalized to a 0-1 scale by subtracting the data by its minimum value and then dividing by its maximum value.

Spindle rotation defects were quantified in two ways. First (Fig. 7E), we scored the number of mCherry::H2B marked chromosomes merged with the cortex at the onset of chromosome separation as described previously (Vargas et al., 2019). Second (Fig. 7F), we recorded the spindle angle relative to the cortex twenty seconds prior to the start of chromosome separation using FIJI (Vargas et al., 2019). Spindles were only analyzed if both poles were present in the same focal plane, indicating a flat orientation, and we recorded the intersecting angle of a line bisecting the spindle in the pole-to-pole axis and a second line tangential to the cortex.

Meiosis I polar body extrusion success (Fig. 7G) was evaluated based on whether oocytes extruded any chromosomes marked by mCherry::H2B into a polar body that remained extruded until meiosis II spindle assembly began, as described previously (Schlientz and Bowerman, 2020).

### Statistics

P-values comparing distributions for all scatter plots were calculated using the Mann–Whitney U-test. P-values comparing slopes were calculated using a two-tailed t-test. Statistical analysis was performed using Microsoft Excel (Microsoft) and Prism 9 (GraphPad Software) and graphs were made in Prism 9.

## SUPPLEMENTAL FIGURE LEGENDS

**FIGURE S1. Meiosis I spindle assembly dynamics in control and TS mutant oocytes at the permissive and restrictive temperatures**. (A-H) Time-lapse maximum intensity projection images during meiosis I in live control and TS mutant oocytes expressing GFP::TBB-2 and mCherry::H2B to mark microtubules and chromosomes, at 15°C (A-D) and at 26°C (E-H). In this and in all subsequent meiosis I time-lapse image series, t=0 is labeled NEBD and is the timepoint immediately preceding the appearance of microtubule bundles forming the cage structure. See the Materials and Methods for a description of the assembly stages and frame selections in this and subsequent supplemental figures. Scale bars = 5 μm.

**FIGURE S2. Timing of temperature shifts in control and TS mutant oocytes**. (A-D) All upshift and downshift timepoints for multipolar and metaphase temperature-shift experiments overlaid onto the timing of meiotic events at 15°C for each allele. Note that the multipolar downshifts are shifted to slightly earlier time points due to the more rapid development at 26°C prior to the downshifts. Error bars and values are mean ± the range. (E) Tables displaying the ranges of multipolar and metaphase temperature-shift timepoints relative to cage onset at 15°C for each allele.

**FIGURE S3. Meiosis I spindle assembly dynamics in control and TS mutant oocytes subjected to prometaphase temperature shifts**. (A-G) Time-lapse maximum intensity projection images of live control and TS mutant oocytes upshifted to 26°C (A-D) or downshifted to 15°C (E-G) during the multipolar stage, in oocytes expressing GFP::TBB-2 and mCherry::H2B. Oocytes depicted in G have cage structure defects (for an example, see Movie S9). Scale bars = 5 μm.

**FIGURE S4. Meiosis I pole assembly dynamics in control and TS mutant oocytes at the permissive and restrictive temperatures**. (A-F) Time-lapse maximum intensity projection images of live control and TS mutant oocytes expressing GFP::ASPM-1 and mCherry::H2B to mark spindle poles and chromosomes, at 15°C (A-C), at 26°C (D-F). Scale bars = 5 μm.

**FIGURE S5. Meiosis I pole assembly dynamics in control and TS mutant oocytes subjected to prometaphase temperature shifts**. (A-F) Time-lapse maximum intensity projection images of live control and TS mutant oocytes expressing GFP::ASPM-1 and mCherry::H2B to mark spindle poles and chromosomes, and upshifted to 26°C (A-C) or downshifted to 15°C (D-F) during the multipolar stage. Montage frames highlight pole coalescence dynamics during the multipolar stage through to the onset of spindle shortening. Scale bars = 5 μm.

**FIGURE S6. Meiosis I pole growth dynamics in control and *tac-1(or455ts)* oocytes subjected to prometaphase temperature shifts**. (M-P) Quantification of control and *tac-1(or455ts)* GFP::ASPM-1 foci volume and intensity (see Materials and Methods) pre- and post-multipolar upshift (A, B) and downshift (C,D). Slopes were compared using a two-tailed t-test to calculate P-values. (E) Table showing the time elapsed pre- and post-multipolar upshift (upper table) and multipolar downshifts (lower table) for control and TS mutants. *, P <0.05; ****, P <0.0001.

**FIGURE S7. Meiosis I pole and chromosome dynamics in control and TS mutant oocytes subjected to metaphase temperature upshifts**. (A-H) Time-lapse maximum intensity projection images of live control and TS mutant oocytes upshifted at metaphase and expressing either GFP::ASPM-1 and mCherry::H2B (A-D) or GFP::TBB-2 and mCherry::H2B (E-H). Montage frames highlight defects following metaphase upshift through to the end of meiosis I. White outlined arrowheads denote ectopic spindle poles and solid white arrowheads indicate chromosome congression errors. The montage in B (top row) is also depicted in Figure 7B. Scale bars = 5 μm.

**FIGURE S8. Imaris snapshots of meiosis I pole and chromosome dynamics in control and TS mutant oocytes subjected to metaphase temperature upshifts**. (A-H) Imaris rotated and snapshot projected time-lapse images (see Materials and Methods) of live TS mutant oocytes upshifted at metaphase expressing GFP::ASPM-1 and mCherry::H2B (A-D) or GFP::TBB-2 and mCherry::H2B (E-H). Montage frames highlight defects following metaphase upshift through to spindle shortening. White outlined arrowheads denote ectopic spindle poles; solid white arrowheads indicate chromosome congression errors. Imaris montages of A-H are of the same oocytes shown in the top rows of Figure S7 as maximum intensity projection montages B-H. Scale bars = 5 μm.

**FIGURE S9. Meiosis I spindle rotation and polar body extrusion in control and TS mutant oocytes subjected to metaphase temperature upshifts**. (A-H) Time-lapse maximum intensity projection images of live control and TS mutant oocytes expressing GFP::ASPM-1 and mCherry::H2B or GFP::TBB-2 and mCherry::H2B upshifted at meiosis I metaphase. Dashed lines depict the oocyte cortex. Montages with a white circle in the last frame indicate failed polar body extrusion. The montage in B (top row) is also depicted in Figure 5B. Montages in E-H are also depicted in Fig. S7E bottom row, Fig. 5I, Fig. S7G middle row, and Fig. S7D bottom row, respectively. (I) Table showing the correlation between spindle rotation defects and polar body extrusion failure. Only oocytes in which both spindle rotation and polar body extrusion could be scored are included. Scale bars = 5 μm.

**FIGURE S10. Meiosis II spindle pole assembly dynamics in control and TS mutant oocytes at the permissive and restrictive temperatures**. (A-H) Time-lapse maximum intensity projection images during meiosis II of live control and TS mutant oocytes expressing GFP::ASPM-1 and mCherry::H2B at 15°C (A-D) and at 26°C (E-H). t=0 is the timepoint when meiosis I chromosome separation ends. Scale bars = 5 μm.

**FIGURE S11. Temperature effects on meiosis I cell cycle timing in control and TS mutant oocytes**. (A-D) The timing of meiotic events in oocytes maintained at 15°C or 26°C, and in oocytes that underwent multipolar and metaphase upshifts, in control and TS mutant oocytes. Error bars and values are mean ± the range. (E) Table showing the mean time to spindle shortening in control and TS mutant oocytes for each temperature condition. Spindle bipolarity and chromosome alignment is rarely achieved in multipolar upshifted or TS mutant oocytes at 26°C and so is not scored; ovulation does occur in TS mutants at 26°C but was not scored.

## SUPPLEMENTAL MOVIE LEGENDS

**Movie S1. Microtubule dynamics in a control oocyte maintained at 26°C throughout meiosis I**. *In utero* time-lapse spinning disk confocal Imaris movie of meiosis I in a control oocyte at 26°C expressing GFP::TBB-2 and mCherry::H2B. For this and all supplemental movies, Z-stacks were collected every 10 seconds; frame rate is 10 frames per second. This oocyte is shown in Figure 1E.

**Movie S2. Microtubule dynamics in a *zyg-9(or623ts)* oocyte maintained at 26°C throughout meiosis I**.

*In utero* time-lapse spinning disk confocal Imaris movie of meiosis I in a *zyg-9(or623ts)* oocyte at 26°C expressing GFP::TBB-2 and mCherry::H2B. Frame rate is 10 frames per second. This oocyte is shown in Figure 1F.

**Movie S3. Microtubule dynamics in a multipolar stage upshifted control oocyte**.

*In utero* time-lapse spinning disk confocal Imaris movie of meiosis I in a control oocyte upshifted at the multipolar stage expressing GFP::TBB-2 and mCherry::H2B. Upshift timepoint is indicated by an orange sphere. Frame rate is 10 frames per second. This oocyte is shown in Figure S3A.

**Movie S4. Microtubule dynamics in a multipolar stage upshifted *zyg-9(or623ts)* oocyte**.

*In utero* time-lapse spinning disk confocal Imaris movie of meiosis I in a *zyg-9(or623ts)* oocyte upshifted at the multipolar stage expressing GFP::TBB-2 and mCherry::H2B. Upshift timepoint is indicated by an orange sphere. Frame rate is 10 frames per second. This oocyte is shown in Figure 2B.

**Movie S5. Microtubule dynamics in a multipolar stage upshifted *zyg-9(or634ts)* oocyte**.

*In utero* time-lapse spinning disk confocal Imaris movie of meiosis I in a *zyg-9(or634ts)* oocyte upshifted at the multipolar stage expressing GFP::TBB-2 and mCherry::H2B. Upshift timepoint is indicated by an orange sphere. Frame rate is 10 frames per second. This oocyte is shown in Figure 2C.

**Movie S6. Microtubule dynamics in a multipolar stage upshifted *tac-1(or455ts)* oocyte**.

*In utero* time-lapse spinning disk confocal Imaris movie of meiosis I in a *tac-1(or455ts)* oocyte upshifted at the multipolar stage expressing GFP::TBB-2 and mCherry::H2B. Upshift timepoint is indicated by an orange sphere. Frame rate is 10 frames per second. This oocyte is shown in Figure 2D.

**Movie S7. Rotation of a control oocyte cage structure**.

*In utero* spinning disk confocal Imaris rotation movie of a normal cage structure restricted to the periphery in a control oocyte at 26°C expressing GFP::TBB-2 and mCherry::H2B.

**Movie S8. Spindle assembly defects in a multipolar stage upshifted *tac-1(or455ts)* oocyte with a normal cage structure**.

*In utero* time-lapse spinning disk confocal Imaris movie of meiosis I in a *tac-1(or455ts)* oocyte upshifted at the multipolar stage expressing GFP::TBB-2 and mCherry::H2B. Rotation displays normal cage structure. Upshift timepoint is indicated by an orange sphere. Frame rate is 10 frames per second. This oocyte is shown in Figure S3D (top row).

**Movie S9. Rescue of bipolar spindle assembly in a multipolar stage downshifted *tac-1(or455ts)* oocyte with a cage structure defect**.

*In utero* time-lapse spinning disk confocal Imaris movie of meiosis I in a *tac-1(or455ts)* oocyte downshifted at the multipolar stage expressing GFP::TBB-2 and mCherry::H2B. Rotation displays abnormal cage structure with microtubule bundles passing in between chromosomes through the internal volume. Downshift timepoint is indicated by a green sphere. Frame rate is 10 frames per second. This oocyte is shown in Figure 2H.

**Movie S10. GFP::ASPM-1 pole splitting event in a multipolar stage upshifted *tac-1(or455ts)* oocyte**.

Excerpt of an *in utero* time-lapse spinning disk confocal Imaris movie of a *tac-1(or455ts)* oocyte upshifted at the multipolar stage expressing GFP::ASPM-1 and mCherry::H2B. Upshift timepoint is indicated by an orange sphere. Frame rate is 3 frames per second.

**Movie S11. Pole assembly dynamics in a multipolar stage upshifted control oocyte**.

*In utero* time-lapse spinning disk confocal Imaris movie of NEBD to spindle shortening onset in a control oocyte upshifted at the multipolar stage expressing GFP::ASPM-1 and mCherry::H2B. Upshift timepoint is indicated by an orange sphere. Frame rate is 5 frames per second. This oocyte is shown in Figure S5A (top row).

**Movie S12. Pole assembly dynamics in a multipolar stage upshifted *zyg-9(or623ts)* oocyte**.

*In utero* time-lapse spinning disk confocal Imaris movie of NEBD to spindle shortening onset in a *zyg-9(or623ts)* oocyte upshifted at the multipolar stage expressing GFP::ASPM-1 and mCherry::H2B. Upshift timepoint is indicated by an orange sphere. Frame rate is 5 frames per second. This oocyte is shown in Figure 4H.

**Movie S13. Pole assembly dynamics in a multipolar stage upshifted *tac-1(or455ts)* oocyte**.

*In utero* time-lapse spinning disk confocal Imaris movie of NEBD to spindle shortening onset in a *tac-1(or455ts)* oocyte upshifted at the multipolar stage expressing GFP::ASPM-1 and mCherry::H2B. Upshift timepoint is indicated by an orange sphere. Frame rate is 5 frames per second. This oocyte is shown in Figure 4I.

**Movie S14. Pole assembly dynamics in a multipolar stage downshifted control oocyte**.

*In utero* time-lapse spinning disk confocal Imaris movie of NEBD to spindle shortening onset in a control oocyte downshifted at the multipolar stage expressing GFP::ASPM-1 and mCherry::H2B. Downshift timepoint is indicated by a green sphere. Frame rate is 5 frames per second. This oocyte is shown in Figure 4J.

**Movie S15. Pole assembly dynamics in a multipolar stage downshifted *zyg-9(or623ts)* oocyte**. *In utero* time-lapse spinning disk confocal Imaris movie of NEBD to spindle shortening onset in a *zyg-9(or623ts)* oocyte downshifted at the multipolar stage expressing GFP::ASPM-1 and mCherry::H2B. Downshift timepoint is indicated by a green sphere. Frame rate is 5 frames per second. This oocyte is shown in Figure 4K.

**Movie S16. Pole assembly dynamics in a multipolar stage downshifted *tac-1(or455ts)* oocyte**. *In utero* time-lapse spinning disk confocal Imaris movie of NEBD to spindle shortening onset in a *tac-1(or455ts)* oocyte downshifted at the multipolar stage expressing GFP::ASPM-1 and mCherry::H2B. Downshift timepoint is indicated by a green sphere. Frame rate is 5 frames per second. This oocyte is shown in Figure 4L.

**Movie S17. Ectopic pole formation in a metaphase stage upshifted *zyg-9(or634ts)* oocyte**.

*In utero* time-lapse spinning disk confocal Imaris movie of a *zyg-9(or634ts)* oocyte upshifted at metaphase expressing GFP::ASPM-1 and mCherry::H2B. Movie is from the point of upshift (indicated by an orange sphere) to the end of meiosis I. Frame rate is 5 frames per second. This oocyte is shown in Figure 5C.

**Movie S18. Chromosome congression defect in a metaphase stage upshifted *zyg-9(or623ts)* oocyte**.

*In utero* time-lapse spinning disk confocal Imaris movie of a *zyg-9(or623ts)* oocyte upshifted at metaphase expressing GFP::ASPM-1 and mCherry::H2B. Movie is from the point of upshift (indicated by an orange sphere) to the end of meiosis I. Frame rate is 5 frames per second. This oocyte is shown in Figure S7B (top row) and Figure 7B.

**Movie S19. Ectopic pole associated with a poorly congressed bivalent in a metaphase stage upshifted *zyg-9(or623ts)* oocyte**.

*In utero* time-lapse spinning disk confocal Imaris movie of a *zyg-9(or623ts)* oocyte upshifted at metaphase expressing GFP::ASPM-1 and mCherry::H2B. Movie is from the point of upshift (indicated by an orange sphere) to the end of meiosis I. Rotation shows the ectopic pole associated with a poorly congressed bivalent. Frame rate is 5 frames per second. This oocyte is shown in Figure S7B (bottom row).

**Movie S20. Ectopic microtubule bundle associated with a poorly congressed bivalent in a metaphase stage upshifted *tac-1(or455ts)* oocyte**.

*In utero* time-lapse spinning disk confocal Imaris movie of a *tac-1(or455ts)* oocyte upshifted at metaphase expressing GFP::TBB-2 and mCherry::H2B. Movie is from the point of upshift (indicated by an orange sphere) to the end of meiosis I. Rotation shows the ectopic microtubule bundle associated with a poorly congressed bivalent. Frame rate is 5 frames per second. This oocyte is shown in Figure 5K.

**Movie S21. Spindle rotation failure in a metaphase stage upshifted *tac-1(or455ts)* oocyte**.

*In utero* time-lapse spinning disk confocal Imaris movie of a *tac-1(or455ts)* oocyte upshifted at metaphase expressing GFP::ASPM-1 and mCherry::H2B. Movie is from the point of upshift (indicated by an orange sphere) to the end of meiosis I. Rotation shows the vertically oriented spindle at point of upshift. This oocyte is shown in Figure 7D.

**Movie S22. Polar body extrusion failure in a metaphase stage upshifted *zyg-9(or634ts)* oocyte**.

*In utero* time-lapse spinning disk confocal Imaris movie of a *zyg-9(or634ts)* oocyte upshifted at metaphase expressing GFP::TBB-2 and mCherry::H2B. Movie is from the point of upshift (indicated by an orange sphere) to the start of meiosis II. Rotation shows both chromosome sets present in the cytoplasm at meiosis II onset. Frame rate is 10 frames per second. This oocyte is shown in Figure S7G (top row) and Figure S9C (bottom row).

**Movie S23. Imaris ‘Spots’ module capturing GFP::ASPM-1 foci in a control oocyte**.

*In utero* time-lapse spinning disk confocal Imaris movie of Spots created on GFP::ASPM-1 signal in a control oocyte upshifted at the multipolar stage expressing GFP::ASPM-1 and mCherry::H2B. This movie is the same as Movie S11 except displaying Imaris Spots. Upshift timepoint is indicated by an orange sphere. Frame rate is 5 frames per second.

## References

Akhmanova, A., and M.O. Steinmetz. 2015. Control of microtubule organization and dynamics: two ends in the limelight. Nat. Rev. Mol. Cell Biol. 16:711–726. doi:10.1038/nrm4084.

Al-Bassam, J., and F. Chang. 2011. Regulation of microtubule dynamics by TOG-domain proteins XMAP215/Dis1 and CLASP. Trends Cell Biol. 21:604–614. doi:10.1016/J.TCB.2011.06.007.

Al-Bassam, J., N.A. Larsen, A.A. Hyman, and S.C. Harrison. 2007. Crystal Structure of a TOG Domain: Conserved Features of XMAP215/Dis1-Family TOG Domains and Implications for Tubulin Binding. Structure. 15:355–362. doi:10.1016/J.STR.2007.01.012.

Bellanger, J.-M., J.C. Carter, J.B. Phillips, C. Canard, B. Bowerman, and P. Gönczy. 2007. ZYG-9, TAC-1 and ZYG-8 together ensure correct microtubule function throughout the cell cycle of C. elegans embryos. J. Cell Sci. 120:2963–73. doi:10.1242/jcs.004812.

Bellanger, J.-M., and P. Gönczy. 2003. TAC-1 and ZYG-9 Form a Complex that Promotes Microtubule Assembly in C. elegans Embryos. Curr. Biol. 13:1488–1498. doi:10.1016/S0960-9822(03)00582-7.

Le Bot, N., M.-C. Tsai, R.K. Andrews, and J. Ahringer. 2003. TAC-1, a Regulator of Microtubule Length in the C. elegans Embryo. Curr. Biol. 13:1499–1505. doi:10.1016/S0960-9822(03)00577-3.

Brenner, S. 1974. THE GENETICS OF CAENORHABDITIS ELEGANS. Genetics. 77:71–94. doi:10.1093/genetics/77.1.71.

van Breugel, M., D. Drechsel, and A. Hyman. 2003. Stu2p, the budding yeast member of the conserved Dis1/XMAP215 family of microtubule-associated proteins is a plus end-binding microtubule destabilizer. J. Cell Biol. 161:359–69. doi:10.1083/jcb.200211097.

Brittle, A.L., and H. Ohkura. 2005. Mini spindles, the XMAP215 homologue, suppresses pausing of interphase microtubules in Drosophila. EMBO J. 24:1387–1396. doi:10.1038/sj.emboj.7600629.

Brouhard, G.J., J.H. Stear, T.L. Noetzel, J. Al-Bassam, K. Kinoshita, S.C. Harrison, J. Howard, and A.A. Hyman. 2008. XMAP215 is a processive microtubule polymerase. Cell. 132:79–88. doi:10.1016/j.cell.2007.11.043.

Chen, L., M. Chuang, T. Koorman, M. Boxem, Y. Jin, and A.D. Chisholm. 2015. Axon injury triggers EFA-6 mediated destabilization of axonal microtubules via TACC and doublecortin like kinase. Elife. 4. doi:10.7554/eLife.08695.

Chuang, C.-H., A.J. Schlientz, J. Yang, and B. Bowerman. 2020. Microtubule assembly and pole coalescence: early steps in Caenorhabditis elegans oocyte meiosis I spindle assembly. Biol. Open. 9. doi:10.1242/BIO.052308.

Connolly, A.A., V. Osterberg, S. Christensen, M. Price, C. Lu, K. Chicas-Cruz, S. Lockery, P.E. Mains, and B. Bowerman. 2014. Caenorhabditis elegans oocyte meiotic spindle pole assembly requires microtubule severing and the calponin homology domain protein ASPM-1. Mol. Biol. Cell. 25:1298–1311. doi:10.1091/mbc.e13-11-0687.

Connolly, A.A., K. Sugioka, C.-H. Chuang, J.B. Lowry, and B. Bowerman. 2015. KLP-7 acts through the Ndc80 complex to limit pole number in C. elegans oocyte meiotic spindle assembly. J. Cell Biol. 210:917–32. doi:10.1083/jcb.201412010.

Cook, B.D., F. Chang, I. Flor-Parra, and J. Al-Bassam. 2019. Microtubule polymerase and processive plus-end tracking functions originate from distinct features within TOG domain arrays. Mol. Biol. Cell. 30:1490–1504. doi:10.1091/mbc.E19-02-0093.

Crowder, M.E., J.R. Flynn, K.P. Mcnally, D.B. Cortes, K.L. Price, P.A. Kuehnert, M.T. Panzica, A. Andaya, J.A. Leary, and F.J. Mcnally. 2015. Dynactin-dependent cortical dynein and spherical spindle shape correlate temporally with meiotic spindle rotation in Caenorhabditis elegans. doi:10.1091/mbc.E15-05-0290.

Cullen, C.F., and H. Ohkura. 2001. Msps protein is localized to acentrosomal poles to ensure bipolarity of Drosophila meiotic spindles. Nat. Cell Biol. 3:637–642. doi:10.1038/35083025.

Danlasky, B.M., M.T. Panzica, K.P. McNally, E. Vargas, C. Bailey, W. Li, T. Gong, E.S. Fishman, X. Jiang, and F.J. McNally. 2020. Evidence for anaphase pulling forces during C. elegans meiosis. J. Cell Biol. 219. doi:10.1083/jcb.202005179.

Davies, T., S.N. Jordan, V. Chand, J.A. Sees, K. Laband, A.X. Carvalho, M. Shirasu-Hiza, D.R. Kovar, J. Dumont, and J.C. Canman. 2014. High-Resolution Temporal Analysis Reveals a Functional Timeline for the Molecular Regulation of Cytokinesis. Dev. Cell. 30:209–223. doi:10.1016/J.DEVCEL.2014.05.009.

Davis-Roca, A.C., N.S. Divekar, R.K. Ng, and S.M. Wignall. 2018. Dynamic SUMO remodeling drives a series of critical events during the meiotic divisions in Caenorhabditis elegans. PLOS Genet. 14:e1007626. doi:10.1371/journal.pgen.1007626.

Dickinson, D.J., and B. Goldstein. 2016. CRISPR-Based Methods for Caenorhabditis elegans Genome Engineering. Genetics. 202:885–901. doi:10.1534/genetics.115.182162.

Divekar, N.S., A.C. Davis-Roca, L. Zhang, A.F. Dernburg, and S.M. Wignall. 2021. A degron-based strategy reveals new insights into Aurora B function in C. elegans. PLOS Genet. 17:e1009567. doi:10.1371/journal.pgen.1009567.

Dumont, J., and A. Desai. 2012. Acentrosomal spindle assembly and chromosome segregation during oocyte meiosis. Trends Cell Biol. 22:241–249. doi:10.1016/J.TCB.2012.02.007.

Fabritius, A.S., M.L. Ellefson, and F.J. McNally. 2011. Nuclear and spindle positioning during oocyte meiosis. Curr. Opin. Cell Biol. 23:78–84. doi:10.1016/J.CEB.2010.07.008.

Gergely, F., D. Kidd, K. Jeffers, J.G. Wakefield, and J.W. Raff. 2000. D-TACC: a novel centrosomal protein required for normal spindle function in the early Drosophila embryo. EMBO J. 19:241–252. doi:10.1093/emboj/19.2.241.

Gigant, E., M. Stefanutti, K. Laband, A. Gluszek-Kustusz, F. Edwards, B. Lacroix, G. Maton, J.C. Canman, J.P.I. Welburn, and J. Dumont. 2017. Inhibition of ectopic microtubule assembly by the kinesin-13 KLP-7 prevents chromosome segregation and cytokinesis defects in oocytes. Development. 144:1674–1686. doi:10.1242/dev.147504.

Gunzelmann, J., D. Rüthnick, T. Lin, W. Zhang, A. Neuner, U. Jäkle, and E. Schiebel. 2018. The microtubule polymerase Stu2 promotes oligomerization of the γ-TuSC for cytoplasmic microtubule nucleation. Elife. 7. doi:10.7554/eLife.39932.

Jordan, S.N., T. Davies, Y. Zhuravlev, J. Dumont, M. Shirasu-Hiza, and J.C. Canman. 2016. Cortical PAR polarity proteins promote robust cytokinesis during asymmetric cell division. J. Cell Biol. 212:39–49. doi:10.1083/jcb.201510063.

Kosco, K.A., C.G. Pearson, P.S. Maddox, P.J. Wang, I.R. Adams, E.D. Salmon, K. Bloom, and T.C. Huffaker. 2001. Control of Microtubule Dynamics by Stu2p Is Essential for Spindle Orientation and Metaphase Chromosome Alignment in Yeast. Mol. Biol. Cell. 12:2870–2880. doi:10.1091/mbc.12.9.2870.

Laband, K., R. Le Borgne, F. Edwards, M. Stefanutti, J.C. Canman, J.-M. Verbavatz, and J. Dumont. 2017. Chromosome segregation occurs by microtubule pushing in oocytes. Nat. Commun. 8:1499. doi:10.1038/s41467-017-01539-8.

Lee, M.J., F. Gergely, K. Jeffers, S.Y. Peak-Chew, and J.W. Raff. 2001. Msps/XMAP215 interacts with the centrosomal protein D-TACC to regulate microtubule behaviour. Nat. Cell Biol. 3:643–649. doi:10.1038/35083033.

Matthews, L.R., P. Carter, D. Thierry-Mieg, and K. Kemphues. 1998. ZYG-9, a Caenorhabditis elegans protein required for microtubule organization and function, is a component of meiotic and mitotic spindle poles. J. Cell Biol. 141:1159–68. doi:10.1083/JCB.141.5.1159.

McNally, K., E. Berg, D.B. Cortes, V. Hernandez, P.E. Mains, and F.J. McNally. 2014. Katanin maintains meiotic metaphase chromosome alignment and spindle structure in vivo and has multiple effects on microtubules in vitro. Mol. Biol. Cell. 25:1037–1049. doi:10.1091/mbc.e13-12-0764.

McNally, K.L., and F.J. McNally. 2005. Fertilization initiates the transition from anaphase I to metaphase II during female meiosis in C. elegans. Dev. Biol. 282:218–230. doi:10.1016/J.YDBIO.2005.03.009.

McNally, K.P., M.T. Panzica, T. Kim, D.B. Cortes, and F.J. McNally. 2016. A novel chromosome segregation mechanism during female meiosis. Mol. Biol. Cell. 27:2576–2589. doi:10.1091/mbc.e16-05-0331.

Mullen, T.J., G. Cavin-Meza, I.D. Wolff, E.R. Czajkowski, N.S. Divekar, J.D. Finkle, and S.M. Wignall. 2022. ZYG-9ch-TOG promotes the stability of acentrosomal poles via regulation of spindle microtubules in C. elegans oocyte meiosis. bioRxiv. 2022.01.04.474888. doi:10.1101/2022.01.04.474888.

Mullen, T.J., A.C. Davis-Roca, and S.M. Wignall. 2019. Spindle assembly and chromosome dynamics during oocyte meiosis. Curr. Opin. Cell Biol. 60:53–59. doi:10.1016/J.CEB.2019.03.014.

O’Rourke, S.M., C. Carter, L. Carter, S.N. Christensen, M.P. Jones, B. Nash, M.H. Price, D.W. Turnbull, A.R. Garner, D.R. Hamill, V.R. Osterberg, R. Lyczak, E.E. Madison, M.H. Nguyen, N.A. Sandberg, N. Sedghi, J.H. Willis, J. Yochem, E.A. Johnson, and B. Bowerman. 2011. A Survey of New Temperature-Sensitive, Embryonic-Lethal Mutations in C. elegans: 24 Alleles of Thirteen Genes. PLoS One. 6:e16644. doi:10.1371/journal.pone.0016644.

Ohkura, H. 2015. Meiosis: an overview of key differences from mitosis. Cold Spring Harb. Perspect. Biol. 7. doi:10.1101/cshperspect.a015859.

Peset, I., J. Seiler, T. Sardon, L.A. Bejarano, S. Rybina, and I. Vernos. 2005. Function and regulation of Maskin, a TACC family protein, in microtubule growth during mitosis. J. Cell Biol. 170:1057–1066. doi:10.1083/jcb.200504037.

Peset, I., and I. Vernos. 2008. The TACC proteins: TACC-ling microtubule dynamics and centrosome function. Trends Cell Biol. 18:379–388. doi:10.1016/J.TCB.2008.06.005.

Pintard, L., and B. Bowerman. 2019. Mitotic Cell Division in Caenorhabditis elegans. Genetics. 211:35–73. doi:10.1534/genetics.118.301367.

Schindelin, J., I. Arganda-Carreras, E. Frise, V. Kaynig, M. Longair, T. Pietzsch, S. Preibisch, C. Rueden, S. Saalfeld, B. Schmid, J.-Y. Tinevez, D.J. White, V. Hartenstein, K. Eliceiri, P. Tomancak, and A. Cardona. 2012. Fiji: an open-source platform for biological-image analysis. Nat. Methods. 9:676–682. doi:10.1038/nmeth.2019.

Schlientz, A.J., and B. Bowerman. 2020. C. elegans CLASP/CLS-2 negatively regulates membrane ingression throughout the oocyte cortex and is required for polar body extrusion. PLOS Genet. 16:e1008751. doi:10.1371/journal.pgen.1008751.

Severson, A.F., G. von Dassow, and B. Bowerman. 2016. Oocyte Meiotic Spindle Assembly and Function. Curr. Top. Dev. Biol. 116:65–98. doi:10.1016/BS.CTDB.2015.11.031.

Severson, A.F., D.R. Hamill, J.C. Carter, J. Schumacher, and B. Bowerman. 2000. The aurora-related kinase AIR-2 recruits ZEN-4/CeMKLP1 to the mitotic spindle at metaphase and is required for cytokinesis. Curr. Biol. 10:1162–71. doi:10.1016/S0960-9822(00)00715-6.

Shirasu-Hiza, M., P. Coughlin, and T. Mitchison. 2003. Identification of XMAP215 as a microtubule-destabilizing factor in Xenopus egg extract by biochemical purification. J. Cell Biol. 161:349–58. doi:10.1083/jcb.200211095.

Sköld, H.N., D.J. Komma, and S.A. Endow. 2005. Assembly pathway of the anastral Drosophila oocyte meiosis I spindle. J. Cell Sci. 118:1745–55. doi:10.1242/jcs.02304.

So, C., K. Menelaou, J. Uraji, K. Harasimov, A.M. Steyer, K.B. Seres, J. Bucevičius, G. Lukinavičius, W. Möbius, C. Sibold, A. Tandler-Schneider, H. Eckel, R. Moltrecht, M. Blayney, K. Elder, and M. Schuh. 2022. Mechanism of spindle pole organization and instability in human oocytes. Science (80-.). 375. doi:10.1126/science.abj3944.

So, C., K.B. Seres, A.M. Steyer, E. Mönnich, D. Clift, A. Pejkovska, W. Möbius, and M. Schuh. 2019. A liquid-like spindle domain promotes acentrosomal spindle assembly in mammalian oocytes. Science. 364:eaat9557. doi:10.1126/science.aat9557.

Srayko, M., A. Kaya, J. Stamford, and A.A. Hyman. 2005. Identification and Characterization of Factors Required for Microtubule Growth and Nucleation in the Early C. elegans Embryo. Dev. Cell. 9:223–236. doi:10.1016/J.DEVCEL.2005.07.003.

Srayko, M., S. Quintin, A. Schwager, and A.A. Hyman. 2003. Caenorhabditis elegans TAC-1 and ZYG-9 Form a Complex that Is Essential for Long Astral and Spindle Microtubules. Curr. Biol. 13:1506–1511. doi:10.1016/S0960-9822(03)00597-9.

Thawani, A., R.S. Kadzik, and S. Petry. 2018. XMAP215 is a microtubule nucleation factor that functions synergistically with the γ-tubulin ring complex. Nat. Cell Biol. 20:575–585. doi:10.1038/s41556-018-0091-6.

Tournebize, R., A. Popov, K. Kinoshita, A.J. Ashford, S. Rybina, A. Pozniakovsky, T.U. Mayer, C.E. Walczak, E. Karsenti, and A.A. Hyman. 2000. Control of microtubule dynamics by the antagonistic activities of XMAP215 and XKCM1 in Xenopus egg extracts. Nat. Cell Biol. 2:13–19. doi:10.1038/71330.

Vargas, E., K.P. McNally, D.B. Cortes, M.T. Panzica, B.M. Danlasky, Q. Li, A.S. Maddox, and F.J. McNally. 2019. Spherical spindle shape promotes perpendicular cortical orientation by preventing isometric cortical pulling on both spindle poles during C. elegans female meiosis. Development. 146. doi:10.1242/DEV.178863.

Vasquez, R.J., D.L. Gard, and L. Cassimeris. 1994. XMAP from Xenopus eggs promotes rapid plus end assembly of microtubules and rapid microtubule polymer turnover. J. Cell Biol. 127:985–93. doi:10.1083/jcb.127.4.985.

Wolff, I.D., M. V. Tran, T.J. Mullen, A.M. Villeneuve, and S.M. Wignall. 2016. Assembly of Caenorhabditis elegans acentrosomal spindles occurs without evident microtubule-organizing centers and requires microtubule sorting by KLP-18/kinesin-12 and MESP-1. Mol. Biol. Cell. 27:3122–3131. doi:10.1091/mbc.e16-05-0291.

Woodruff, J.B., B. Ferreira Gomes, P.O. Widlund, J. Mahamid, A. Honigmann, and A.A. Hyman. 2017. The Centrosome Is a Selective Condensate that Nucleates Microtubules by Concentrating Tubulin. Cell. 169:1066–1077.e10. doi:10.1016/J.CELL.2017.05.028.

Yang, H., P.E. Mains, and F.J. McNally. 2005. Kinesin-1 mediates translocation of the meiotic spindle to the oocyte cortex through KCA-1, a novel cargo adapter. J. Cell Biol. 169:447–457. doi:10.1083/jcb.200411132.

Yang, H., K. McNally, and F.J. McNally. 2003. MEI-1/katanin is required for translocation of the meiosis I spindle to the oocyte cortex in C. elegans☆. Dev. Biol. 260:245–259. doi:10.1016/S0012-1606(03)00216-1.

Zhang, L., J.D. Ward, Z. Cheng, and A.F. Dernburg. 2015. The auxin-inducible degradation (AID) system enables versatile conditional protein depletion in C. elegans. Development. 142:4374–84. doi:10.1242/dev.129635.

